# Transcriptomic and epigenomic consequences of heterozygous loss of function mutations in *AKAP11*, the first large-effect shared risk gene for bipolar disorder and schizophrenia

**DOI:** 10.1101/2024.03.14.584883

**Authors:** Nargess Farhangdoost, Calwing Liao, Yumin Liu, Martin Alda, Patrick A. Dion, Guy A. Rouleau, Anouar Khayachi, Boris Chaumette

## Abstract

The gene A-kinase anchoring protein 11 (*AKAP11*) recently emerged as a shared risk factor between bipolar disorder and schizophrenia, driven by large-effect loss-of-function (LoF) variants. Recent research has uncovered the neurophysiological characteristics and synapse proteomics profile of *Akap11*-mutant mouse models. Considering the role of AKAP11 in binding cAMP-dependent protein kinase A (PKA) and mediating phosphorylation of numerous substrates, such as transcription factors and epigenetic regulators, and given that chromatin alterations have been implicated in the brains of patients with bipolar disorder and schizophrenia, it is crucial to uncover the transcriptomic and chromatin dysregulations following the heterozygous knockout of *AKAP11*, particularly in human neurons. In this study, we use genome-wide approaches to investigate such aberrations in human induced pluripotent stem cell (iPSC)-derived neurons. We show the impact of heterozygous *AKAP11* LoF mutations on the gene expression landscape and profile the methylomic and acetylomic modifications. Altogether we highlight the involvement of aberrant activity of intergenic and intronic enhancers, which are enriched in PBX homeobox 2 (PBX2) and Nuclear Factor-1 (NF1) known binding motifs, respectively, in transcription dysregulations of genes functioning as DNA-binding transcription factors, actin and cytoskeleton regulators, and cytokine receptors, as well as genes involved in G-protein-coupled receptors (GPCRs) binding and signaling. A better understanding of the dysregulations resulting from haploinsufficiency in *AKAP11* improves our knowledge of the biological roots and pathophysiology of BD and SCZ, paving the way for better therapeutic approaches.

## Introduction

Bipolar disorder (BD) is a complex, severe psychiatric disorder with a lifetime prevalence of 2-3% in the general population^1,2,4,5^. BD is characterized by recurrent episodes of mania and depression and is often accompanied by psychotic symptoms, resembling those observed in schizophrenia (SCZ)^3,4^, and considerable clinical comorbidities ^1-5^. Despite extensive research, the etiology of BD remains largely unknown. Recent studies have underscored the significance of protein-truncating variations (PTVs) and copy number variations (CNVs) in BD^6,7^. Most recently, the Bipolar Exome (BipEx) consortium, involving whole-exome sequencing of 13,933 BD cases and 14,422 controls, identified *AKAP11* as a significant rare-variant risk gene for BD^3^. *AKAP11* was found to be enriched in rare protein-truncating variations, significantly elevating the risk for BD and establishing it as the strongest known risk gene for BD to date^3^. Notably, this genetic risk factor is shared between BD and SCZ (combined odds ratio [OR] = 7.06, p = 2.83 × 10E−9)^3^.

Generally, A-kinase anchoring proteins (AKAPs) bind to PKA regulatory subunits as well as cytoskeleton proteins or organelles^8,9^, which allows for their pivotal role in distributing and localizing PKA to its targets within the cells^8,9^. They are now understood to have multiple binding sites for various signaling molecules, such as protein kinase C, protein phosphatases, GPCRs, adenylyl cyclases, and phosphodiesterases^10-13^. In neurons, AKAPs localize these signaling enzymes to their substrates allowing precise temporal and spatial interactions among postsynaptic signaling molecules, impacting synaptic plasticity and neuronal function^10-13^. AKAP11 is a ubiquitously expressed membrane-associated and vesicular anchoring protein that is under LoF constraint (LOEUF = 0.32, pLI =1 on gnomAD v4.0) and is highly expressed in the brain tissue^14^. It binds to type II regulatory subunits of PKA and regulates the interaction between PKA and its numerous substrates^15,16^. AKAP11’s interaction with PKA leads to the localization of this complex to its substrates, e.g., Glycogen Synthase Kinase 3 Beta (GSK3B^17^, a molecular target of lithium which is widely used to treat BD^18-21^), IQ motif-containing GTPase-activating protein 2 (IQGAP2)^22^, and protein phosphatase-1 (PP1)^17,23^, regulating their functions. AKAP11-mediated phosphorylation of substrates through PKA can affect a plethora of events, including signaling pathways, regulation of transcription factors, e.g., cAMP-response element binding (CREB) ^24^, and modulation of epigenetic regulators, e.g., histone deacetylases (HDACs)^25,26^. In addition, AKAP11 has been suggested to interact with other proteins including transcription factors and epigenetics regulators, e.g., SET and MYND domain containing 2 (SMYD2/KMT3C)^27-29^ and Iroquois Homeobox 3 (IRX3)^30^. Owing to such potential direct or indirect effects of AKAP11 on the regulation of chromatin accessibility and gene expression, it is important to understand the transcriptomic and epigenomic consequences of heterozygous LoF of *AKAP11* and identify the key processes that overlap with our current knowledge of the etiology of BD and SCZ.

Recently, heterozygous *Akap11* mouse mutants were generated and screened for identification of a neurophysiological biomarker, i.e. electroencephalography (EEG) phenotypes similar to those observed in human SCZ patients were detected^31-33^. Another study carried out a deep proteomics analysis of synapses in *Akap11*-mutant mice and detected some overlap between the dysregulated pathways in SCZ and BD patients and the mutant model^34^. However, our study is the first to investigate the effect of heterozygous LoF mutations in *AKAP11* at the transcriptomic and epigenetic (DNA methylation and histone modification) levels in a human cellular model. A better understanding of the dysregulations resulting from AKAP11 haploinsufficiency may improve our knowledge of the biological roots and pathophysiology of BD and SCZ, eventually contributing to improved diagnosis and therapeutic approaches.

To investigate the transcriptomic and epigenetic consequences of heterozygous LoF of AKAP11, we employed a human iPSC, reprogrammed from lymphoblastoid cells derived from an unaffected individual^35,36^. Subsequently, three heterozygous *AKAP11*-Knockout (*AKAP11-*KO) isogenic clones were created from the iPSC, each harboring a different heterozygous frameshift mutation using CRISPR-Cas9 genome editing. We then differentiated these into 25-day-old neurons and compared their respective transcriptome to the one of WT counterpart neurons. Gene set enrichment analysis (GSEA) highlighted downregulation of genes related to ribosome function and DNA-binding transcription factor activity, and upregulation in molecular functions including signaling receptor interactions, cytokine activity, and cytoskeletal dynamics. Using whole-genome bisulfite sequencing (WGBS) we identified the differentially methylated regions (DMRs) and found an overlap between the genes associated with DMRs and genes with differential expression. These overlapping genes were enriched in GO terms related to DNA-binding transcription factor activity. Examination of enhancer activity in heterozygous *AKAP11*-KO iPSC-derived neurons, using chromatin immunoprecipitation followed by sequencing (ChIP-seq) of histone H3 Lysine 27 acetylation (H3K27ac) mark, revealed that reduced intergenic enhancer activity play a partial role in downregulation of neighboring genes involved in transcription factor binding and activity. Conversely, elevated intronic enhancer activity partly accounts for the upregulation of genes involved in signaling receptor binding, cytokine receptor binding, and cytoskeleton-related functions. Furthermore, we found significant enrichment of GPCR binding and signaling terms among differentially expressed genes (DEGs) in *AKAP11*-KO, as well as among genes associated with DMRs. While DMRs did not seem to particularly influence the transcription of genes involved in such pathways, a portion of such gene expression dysregulations was linked to enhancer activity alterations. Lastly, we investigated the correlation between the DMRs and enhancer activity alterations, marked by H3K27ac, and subsequently pinpointed the most profoundly impacted intergenic and intronic regulatory regions linked to concordant gene expression dysregulations, resulting from heterozygous LoF mutations in *AKAP11*.

## Materials / Subjects and Methods

### Ethics statement

The cell line and protocols in the present study were used in accordance with guidelines approved by the institutional human ethics committee guidelines and the Nova Scotia Health Authority Research Ethics Board (REB # 1020604).

### RNP-CRISPR-Cas9 knockout of *AKAP11* in induced pluripotent stem cells

The iPSCs used in this study were reprogrammed from the lymphoblasts (blood sample) of a consenting individual ^35,36^ (Table 1). We utilized a control iPSC line (SBP009, male, age 53, Caucasian; Table 1) to knock out *AKAP11*, using the RNP-CRISPR-Cas9 genome editing technique. To generate three isogenic knockout clones from the WT iPSC (SBP009), Ribonucleoprotein (RNP)-mediated CRISPR-Cas9 genome editing was performed using the Alt-R CRISPR-Cas9 system (IDT). Two synthetic crRNA guides were designed (TACTGGTATAAGGTACACCT PAM:TGG and CTGATGCAAGAATGTGTCCA PAM:AGG) to form a duplex with Alt-R® CRISPR-Cas9 tracrRNA, ATTO 550 (IDT, Cat. # 1075928) and coupled to the Alt-R© S.p. Cas9 Nuclease V3 (IDT). The Lipofectamine™ Stem Transfection (Invitrogen, Cat. #STEM00001) and PLUS™ Reagent (Invitrogen, Cat. #11514015) ratios were optimized with a lower volume than the manufacturer’s protocol. The transfected cells were incubated for 48 hours. Single ATTO550+ cells were then sorted into 96-well plates. Clones were expanded and individually verified with MiSeq sequencing of the target loci using the following primers: fwLeft: CCAAGGAGCTTTTCAGGATGC and fwRight: TGGTCTCAAGTTGGTGCTTCT. All the clones were also kept frozen at the NPC stage in FBS (Gibco, Cat. #12484028) +10% DMSO (Sigma, Cat. #D2650) at the lowest passage possible for future use. The control group consists of three replicates of SBP009 iPSCs at different passage numbers that were separately induced into NPCs, through separate EB formation and rosette selection processes. These NPCs were then differentiated into 25-day-old neurons using the same protocol, as described below.

**Table 1.**
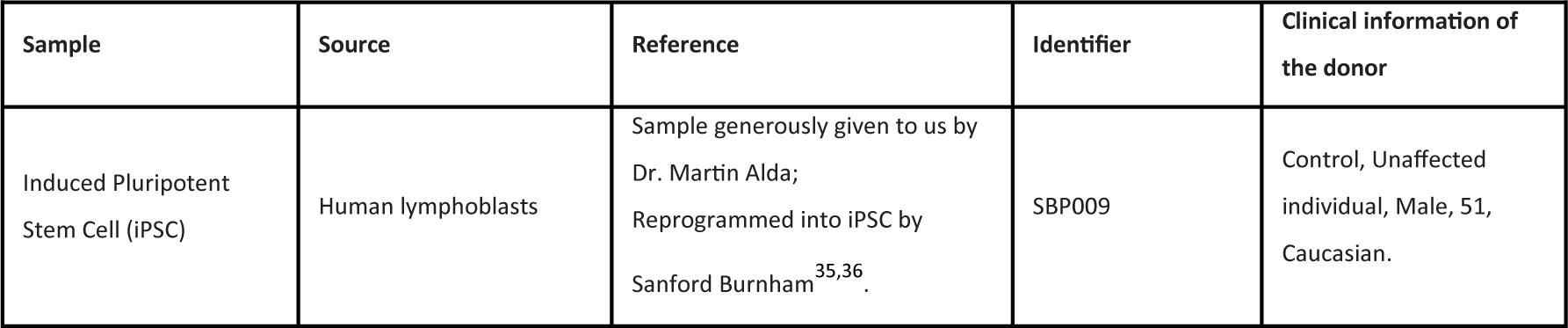
Details of the iPSC sample used in the study.

### iPSC and NPC differentiation and maintenance and neuronal cell culture

iPSCs were maintained on a Matrigel (Corning, Cat. #08-774-552) coated plate with mTeSR™1 medium (STEMCELL Technologies, Cat. #85870) along with Y-27632 (ROCK inhibitor, STEMCELL Technologies, Cat. # 72307) after each passage. During the course of induction of iPSC to NPCs (20 days), STEMdiff™ SMADi Neural Induction Kit (STEMCELL Technologies, Cat. #08581) was used. The method for induction of iPSC to NPCs was based on the STEMCELL™ neural induction EB-based method and was performed following the manufacturer’s protocol. On Day 0, a single-cell suspension of iPSC was generated using Gentle Cell Dissociation Reagent (STEMCELL Technologies, Cat. # 100-0485), and 10,000 cells per microwell were seeded in an ultralow attachment 96-well plate to generate Embryoid bodies (EBs), cultured in STEMdiff™ Neural Induction Medium + SMADi (STEMdiff™ SMADi Neural Induction Kit, Cat. # 08582) + 10 μM Y-27632. From Day 1 to Day 4, a daily partial (3/4) medium change was performed using STEMdiff™ Neural Induction Medium + SMADi. On Day 5, EBs were harvested using a wild-board 1ml serological pipettes and 40μm strainer and transferred to a single well of a 6-well plate coated with Poly-L-ornithine hydrobromide (PLO) and laminin. A daily full medium change was performed from Day 6 to Day 11. After the neural induction efficiency was determined higher than 75%, neural rosettes were manually selected using STEMdiff™ Neural Rosette Selection Reagent (STEMCELL Technologies, Cat. # 05832) on Day 12 and replated onto a single well of PLO/Laminin coated 6-well plate. With continuous daily full medium change, selected rosette-containing clusters attached, and NPC outgrowths formed a monolayer between the clusters. The NPCs were ready for passage 1 when cultures were approximately 80 - 90% confluent (typically on Day 19). NPCs were maintained and expanded (7-10 days) using DMEM/F12 with Glutamax (ThermoFisher, Cat. #10565042), N2 1x (ThermoFisher, Cat. # 17502001), B27 1x (ThermoFisher, Cat. #17504001), FGF2 20 ng/ml (PEPROTECH, Cat. #100-18B), and EGF 20 ng/ml (Gibco, Cat. # AF-100-15-1MG). For the final differentiation of NPCs into neurons, NPCs were detached using Accutase (STEMCELL Technologies, Cat. #7922) and then differentiated onto PLO/laminin-coated plates with final differentiation media of BrainPhys™ Neuronal Medium (STEMCELL Technologies, Cat. #05790) supplemented with Glutamax 1x (ThermoFisher, Cat. #35050061), N2 1x (ThermoFisher, Cat. #17502001), B27 1x (ThermoFisher, Cat. #17504001), 200nM ascorbic acid 200nM (STEMCELL Technologies, Cat. #72132), cyclic AMP 500 μg/ml (Sigma-Aldrich, Cat #A9501-1G), brain-derived neurotrophic factor 20 ng/ml (BDNF, GIBCO, Cat. #PHC7074), Wnt3a 10 ng/ml (R&D Systems, Cat. #5036-WN-500), and laminin 1 μg/ml (Gibco, Cat. #23017015) for 14 days (medium change of 3 times per week). Subsequently, the cells were maintained in STEMdiff™ Forebrain Neuron Maturation Kit for 11 days (STEMCELL Technologies, Cat. #08605) --final differentiation media was replaced by STEMdiff™ Forebrain Neuron Maturation Kit in the same dishware.

### Reverse-transcription quantitative PCR (RT-qPCR)

To confirm the lower *AKAP11* expression in KOs compared to the parental, reverse-transcription quantitative PCR (RT-qPCR) was performed on neurons using a pre-design assay from ThermoScientific (Cat. #4351372, Assay ID: Hs01568657_g1, Ref gene: *POLR2A*) targeting *AKAP11* exon 7, where the CRISPR-Cas9 guide sites were located. RNA extraction was performed using RNeasy kit Qiagen (Cat. #74004) according to the manufacturer’s instructions.

### Neuronal immunocytochemistry

iPSC-derived neurons were fixed in phosphate-buffered saline (PBS) containing 5% sucrose and 3.7% formaldehyde for 1 h at room temperature (RT). They were then permeabilized with the same buffer but with 0.2% Triton 100x for 2 minutes at RT. The cells were rinsed twice with PBS 1x at RT and incubated with NH4CI (50mM) in PBS for 10 minutes at RT. They were then rinsed with PBS 1x twice. It was ensured that the coverslip did not dry out. The cells were incubated in PBS1x / goat serum (GS) 10% for 20 minutes at RT. Primary antibody (BIII-Tubulin from Sigma-Aldrich (mouse, Cat. #T8660)) along with DAPI (Abcam; Cat. #ab228549) were added to the cells in PBS 1x / 5% GS / 0.05% Triton 100x and were incubated overnight at 4°C. The cells were then washed three times (5 minutes each) with PBS 1x at RT. The appropriate secondary antibody (1/1000) conjugated to Alexa488 (Invitrogen, Cat. #A21202) was added to the cells in PBS 1x / 5% GS / 0.05% Triton 100x and incubated for 45 minutes at RT. The cells were then washed three times with PBS at RT and once with ddH2O. Finally, the coverslip was mounted with Prolong (ThermoFisher, Cat. #P36930) until confocal examination. The images (1024 × 1024) were acquired with a ×20 lens (numerical aperture NA 1.4) on an SP8 confocal microscope (Leica Microsystems) and were visualized using image j (FIJI).

### RNA-seq library Preparation and Sequencing

RNA was extracted from bulk neuronal culture 25 days post-differentiation using RNeasy Mini Kit (Cat. #74104). Library preparation from total RNA was performed using NEB rRNA-depleted (HMR) stranded library preparation kit according to the manufacturer’s instructions and sequencing was carried out using Illumina NovaSeq 6000 (100 bp paired-end).

### WGBS library preparation and sequencing

DNA was extracted from bulk neuronal culture 25 days post-differentiation using DNeasy Blood & Tissue Kit (Cat. #69504). Whole genome sequencing libraries were generated from 1000 ng of genomic DNA spiked with 0.1% (w/w) unmethylated λ DNA (Roche Diagnostics) and fragmented to 300–400 bp peak sizes using the Covaris focused-ultrasonicator E210. Fragment size was controlled on a Bioanalyzer High Sensitivity DNA Chip (Agilent) and NxSeq AmpFREE Low DNA Library Kit (Lucigen) was applied. End repair of the generated dsDNA with 3′ or 5′ overhangs, adenylation of 3′ ends, adaptor ligation, and clean-up steps were carried out as per Lucigen’s recommendations. The cleaned-up ligation product was then analyzed using LabChip GX/GX II (Caliper). Samples were then bisulfite converted using the EZ-DNA Methylation Gold Kit (Zymo Research) according to the manufacturer’s protocol. DNA was amplified by 9 cycles of PCR using the Kapa HiFi Uracil+ Kit (Roche) DNA polymerase (KAPA Biosystems) according to the manufacturer’s protocol. The amplified libraries were purified using Ampure XP Beads (Beckman Coulter), validated on LabChip GX/GX II (Caliper), and quantified by qPCR amplification with Illumina adapters. The values are read using Lightcycler LC480 (Roche). Sequencing of the WGBS libraries was performed on the Illumina NovaSeq6000 S4 v1.5 using 150-bp paired-end sequencing.

### Cross-linking, ChIP-seq library preparation, and sequencing

About 10 million neuronal cells per sample (3 heterozygous *AKAP11*-KO clones and 3 WT replicates) were grown and directly crosslinked on the plate with 1% formaldehyde (Sigma) for 10 minutes at room temperature and the reaction was stopped using 125nM Glycine for 5 minutes. Fixed cell preparations were washed with ice-cold PBS, scraped off the plate, pelleted, washed twice again in ice-cold PBS, and the flash-frozen pellets were stored at −80°C.

Thawed pellets were resuspended in 500ul cell lysis buffer (5 mM PIPES-pH 8.5, 85 mM KCl, 1% (v/v) IGEPAL CA-630, 50 mM NaF, 1 mM PMSF, 1 mM Phenylarsine Oxide, 5 mM Sodium Orthovanadate, EDTA-free Protease Inhibitor tablet) and incubated 30 minutes on ice. Samples were centrifugated and pellets resuspended in 500ul of nuclei lysis buffer (50 mM Tris-HCl pH 8.0, 10 mM EDTA, 1% (w/v) SDS, 50 mM NaF, 1 mM PMSF, 1 mM Phenylarsine Oxide, 5 mM Sodium Orthovanadate and EDTA-free protease inhibitor tablet) and incubated 30 minutes on ice. Sonication of lysed nuclei was performed on a BioRuptor UCD-300 at max intensity for 45 cycles, 10 s on 20 s off, centrifuged every 15 cycles, and chilled by a 4°C water cooler. Samples were checked for sonication efficiency using the criteria of 150–500bp by gel electrophoresis of a reversed cross-linked and purified aliquot. After the sonication, the chromatin was diluted to reduce the SDS level to 0.1% and concentrated using Nanosep 10k OMEGA (Pall). ChIP reaction for histone modifications was performed on a Diagenode SX-8G IP-Star Compact using Diagenode automated Ideal ChIP-seq Kit for Histones. Dynabeads Protein A (Invitrogen) were washed, then incubated with specific antibodies (rabbit polyclonal anti-H3K27ac Diagenode Cat. #C15410196, RRID: AB_2637079), 1 million cells of sonicated cell lysate, and protease inhibitors for 10 hr, followed by 20 min wash cycle using the provided wash buffers (Diagenode Immunoprecipitation Buffers, iDeal ChIP-seq kit for Histone).

Reverse cross-linking took place on a heat block at 65°C for 4 hr. ChIP samples were then treated with 2ul RNase Cocktail at 65°C for 30 min followed by 2ul Proteinase K at 65°C for 30 min. Samples were then purified with QIAGEN MinElute PCR purification kit (QIAGEN) as per manufacturers’ protocol. In parallel, input samples (chromatin from about 50,000 cells) were reverse crosslinked, and DNA was isolated following the same protocol. Library preparation was carried out using Kapa Hyper Prep library preparation reagents (Kapa Hyper Prep kit, Roche 07962363001) following the manufacturer’s protocol. ChIP libraries were sequenced using Illumina NovaSeq 6000 at 100bp paired-end reads.

### RNA-seq data processing and analysis

Adaptor sequences and low-quality score bases (Phred score < 30) were first trimmed using Trimmomatic^37^. The resulting reads were aligned to the GRCh38 human reference genome assembly, using STAR^38^. Read counts were obtained using HTSeq^39^ with parameters-m intersection-nonempty-stranded=reverse. For all downstream analyses, we excluded lowly expressed genes with an average read count lower than 20 across all samples, resulting in a total of 23,590 genes. Raw counts were normalized using edgeR’s TMM algorithm^40^ and were then transformed to log2-counts per million (logCPM) using the voom function implemented in the limma R package^41^.

To assess differences in gene expression levels between the different conditions (3 KOs versus 3 WT replicates), we fitted a linear model using limma’s lmfit function with parameter method="robust". Nominal p-values were corrected for multiple testing using the Benjamini-Hochberg method.

The Z-score heatmap was constructed using the R package ComplexHeatmap^42^; KO vs WT analysis: P-adj < 0.05 & | log2FC| > 1.

Gene set enrichment analysis (GSEA) based on a pre-ranked gene list by t-statistic was performed using the R package fgsea (http://bioconductor.org/packages/fgsea), P-adj < 0.05. Over-representation analysis (ORA) was performed using ShinyGO 0.77^43^ (http://bioinformatics.sdstate.edu/go) and FDR<0.05. The background genes used for ORA were all the 23,590 genes that were considered expressed. FDR is calculated based on the nominal P-value from the hypergeometric test. Fold Enrichment is the percentage of genes in the input gene list belonging to a pathway divided by the corresponding percentage in the background.

### For visualization, gene expression data was corrected for batch effects using ComBat^44^. WGBS data processing and analysis

Adaptor sequences and low-quality score bases were first trimmed using Trimmomatic^37^. The resulting reads were mapped to the human reference genome (GRCh38) and lambda phage genome using Bismark^45^, which uses a bisulfite-converted reference genome for read mapping. Only reads that had a unique alignment were retained and PCR duplicates were marked using Picard tools (https://broadinstitute.github.io/picard/). Methylation levels for each CpG site were estimated by counting the number of sequenced C (‘methylated’ reads) divided by the total number of reported C and T (‘unmethylated’ reads) at the same position of the reference genome using Bismark’s methylation extractor tool. We performed a strand-independent analysis of CpG methylation where counts from the two Cs in a CpG and its reverse complement (position on the plus strand and position i+1 on the minus strand) were combined and assigned to the position of the C in the plus strand. To assess MethylC-seq bisulfite conversion rate, the frequency of unconverted cytosines (C basecalls) at lambda phage CpG reference positions was calculated from reads uniquely mapped to the lambda phage reference genome. CpG sites with at least 5X coverage were included for all downstream analyses. Differential methylation analyses were performed using the R (R 4.1.2, https://www.r-project.org/) package DSS (DMLfit.multiFactor function with parameter smoothing = TRUE)^46^. DMRs were called using the function callDMR at default parameters with a minimum length of 50 base pairs and 3 CpG sites (delta=0, p.threshold=1e-5, minlen=50, minCG=3, dis.merge=100, pct.sig=0.5). Annotation determined using Homer (hg38)^47^. Methylation values and P-values are averaged within each DMR.

Over-representation analysis of genes assigned to DMRs was performed using ShinyGO 0.77^43^ (http://bioinformatics.sdstate.edu/go) with FDR>0.05. The background genes used for ORA were the same background gene list (23,590 genes) obtained from our RNA-seq analysis. FDR is calculated based on the nominal P-value from the hypergeometric test. Fold Enrichment is the percentage of genes in the input gene list belonging to a pathway divided by the corresponding percentage in the background.

### ChIP-seq data processing and analysis

ChIP-seq reads were first trimmed for adapter sequences and low-quality score bases using Trimmomatic^37^. The resulting reads were mapped to the human reference genome (GRCh38) using BWA-MEM^48^ in paired-end mode at default parameters. Only reads that had a unique alignment (mapping quality > 20) were retained and PCR duplicates were marked using Picard tools (http://broadinstitute.github.io/picard/). Peaks were called using MACS2^49^.

To detect changes in histone modifications, a consensus peak set for H3K27ac was first generated by merging ChIP-seq peaks across samples using bedtools merge (https://bedtools.readthedocs.io/). A peak must be present in at least one sample in either condition. Read counts were obtained within these genomic regions using HOMER. Differential peak analysis was performed using the R package limma^41^. Nominal p-values were corrected for multiple testing using the Benjamini-Hochberg method.

Peaks were associated with the nearest TSS of genes using the annotatePeaks command from HOMER software suite ^47^. Enrichment analysis (ORA) of genes assigned to differential peaks was performed using ShinyGO 0.77 ^43^ (http://bioinformatics.sdstate.edu/go). The background genes used for ORA were the same background gene list (23,590 genes) obtained from our RNA-seq analysis. FDR is calculated based on the nominal P-value from the hypergeometric test. Fold Enrichment is the percentage of genes in the input gene list belonging to a pathway divided by the corresponding percentage in the background. Motif enrichment within differential peaks was performed using the HOMER findMotifsGenome command. We only included genes/peaks that are associated within 100 kb (distance of 100 kb from the TSS to the peak center) and have concordant changes in expression/histone mark.

Bigwig coverage files were created with the HOMER makeUCSCfile command and bedGraphToBigWig utility from UCSC. Data were normalized so that each value represents the read count per base pair per 10 million reads. Heatmaps and average profiles were generated using modules “computeMatrix” (--referencePoint center) followed by “plotHeatmap” and “plotProfile” from deepTools^50^, using bigwig files as input.

## Results

### Heterozygous *AKAP11* knockout in human iPSCs and generation of iPSC-derived neurons

Using a human iPSC, from the lymphoblasts of an unaffected individual^35,36^, we generated 3 heterozygous *AKAP11*-knockout (*AKAP11-*KO) isogenic neuronal clones, each carrying a different heterozygous frameshift mutation (Figure 1A). We then validated the heterozygous *AKAP11-*KO isogenic clones (at the iPSC level) using Miseq amplicon sequencing. Among the individual clones obtained, clone #16 has an 11 bp-deletion, clone #20 has a 10 bp-deletion, and clone #21 has a 4 bp-insertion (Supplementary Figure 1A). Additionally, we performed a KaryoStat+ assay for digitally visualizing any probable chromosomal aberrations in the genetically edited iPSC clones, and we confirmed that there are no chromosomal aberrations in the isogenic heterozygous *AKAP11-*KO iPSC clones compared to their WT counterpart (Supplemental Figure 1B). Subsequently, the *3* heterozygous *AKAP11-*KO iPSC clones, along with 3 replicates of their parental iPSC counterparts (WT; SBP009) were induced into neural progenitor cells (NPCs) and were then differentiated into forebrain neurons of age 25 days (Figure 1A; both groups of neurons were collected on day 25). After collecting the neurons, we carried out RT-qPCR to ensure the gene expression level changes of the heterozygous *AKAP11-*KO clones compared to the WT counterpart in the neuronal context and observed significantly lower expression levels of the heterozygous *AKAP11-*KO clones compared to the WT parental. Clones 16 and 21 had a 0.4-fold and clone 20 had a 0.6-fold decrease in *AKAP11* expression of exon 7 (Figure 1B). We then performed Neuronal Immunocytochemistry and stained the heterozygous *AKAP11-*KO and WT neurons with BIII-Tubulin, a neuronal marker, and DAPI and confirmed the presence of a viable neuronal culture broadly expressing BIII-Tubulin (Figure 1C). This ensures that the heterozygous knockout of *AKAP11* does not disturb the neuronal culture differentiation and viability. Further cell type deconvolution using bulk RNA sequencing (RNA-seq) data and dtangle method^51^ confirmed the presence of neurons as the most abundant cell type in our cultures for both heterozygous *AKAP11*-KO and WT samples (Supplementary Figure 1C).

**Figure 1.**
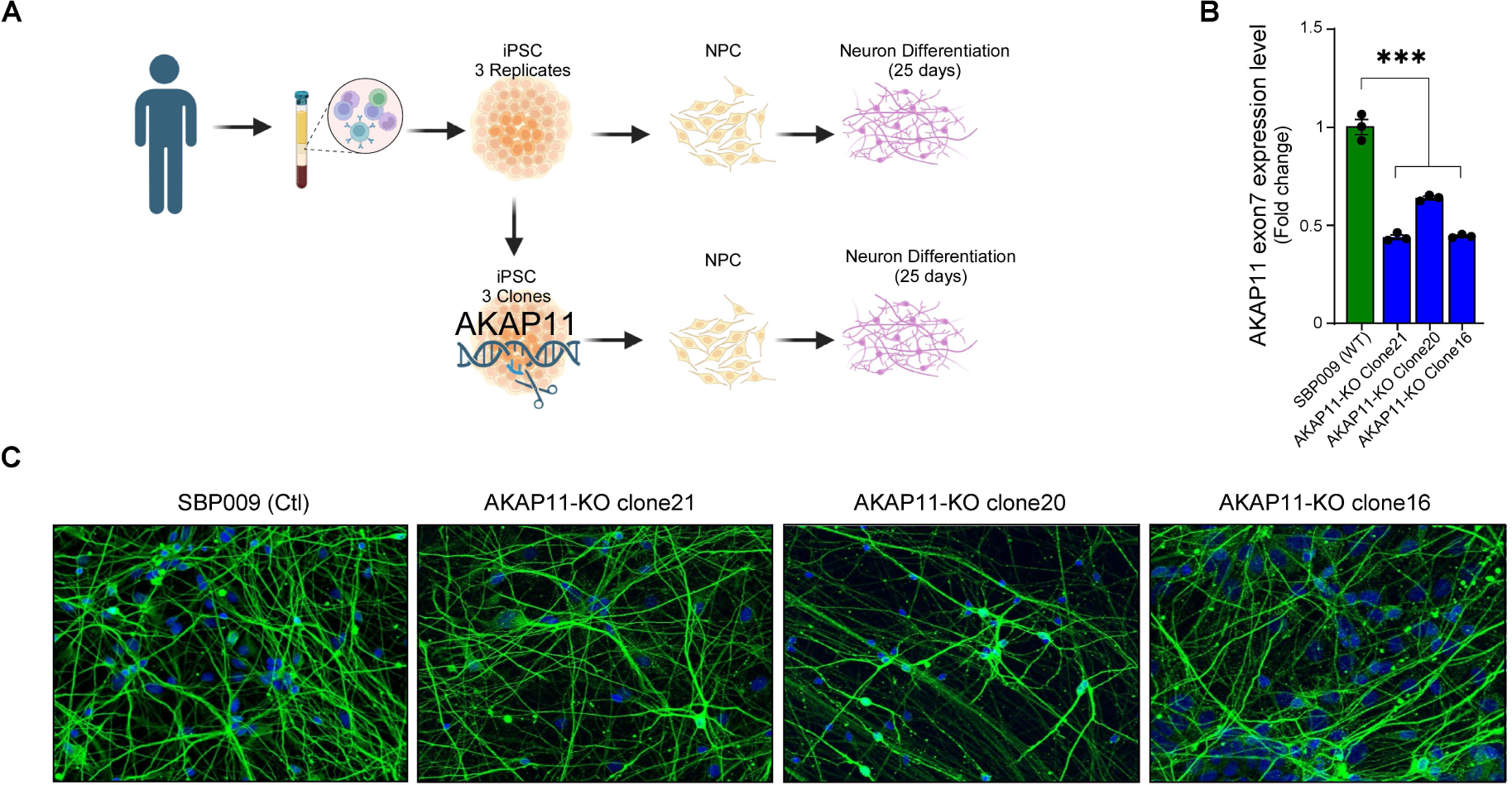
iPSC differentiation into neurons, validation of *AKAP11*-KO, and culture viability. **A**) Schematic view of the origin and differentiation steps of iPSC-derive neurons as well as heterozygous knockout of *AKAP11* at iPSC level. **B**) R T-qPCR gene expression level changes of exon 7 of the *AKAP11*-KO clones and WT (SBP009). Each clone was processed as triplicates along with an endogenous control (POLR2A). Data shown in this figure are the mean ± s.e.m. and statistical significance determined by one-way ANOVA with Bonferroni post-test. ****p<0.0001. **C**) Neuronal Immunocytochemistry of *AKAP11*-KO iPSC-derived neuronal clones at day 25 post-differentiation. BIII-Tubulin: Green, DAPI: Blue.

### Transcriptomic changes and biological pathways affected by heterozygous LoF of *AKAP11* in iPSC-derived neurons

Using the gene expression data from RNA-seq, we identified DEGs resulting from heterozygous *AKAP11-*KO vs. WT comparison with the p-adjusted value of (p-adj) < 0.05 and |log2FC| > 1 (Figure 2A, Supplementary Table 1). A total of 849 genes showed significant differential expression out of which 421 genes were downregulated, and 428 were upregulated (Figure 2B). Hierarchical clustering of differentially expressed genes demonstrated a clear separation in gene expression profiles between heterozygous *AKAP11*-KO and WT with the data pattern similarity being high within groups but low between groups (Figure 2C). The top 30 downregulated DEGs, ranked by their p-values, primarily include coding genes and long-noncoding RNA (lncRNA) genes, four of which are located in Prader-Willi syndrome (PWS) region (*PWRN1*, *PWRN4*, *NDN*, *MAGEL2*), and several are related to the homeobox (HOX) gene family (*HOTAIRM1*, *PHOX2A*, *DRGX*, *HOXA2*, *HOXC8*). The top 30 upregulated DEGs include coding and lncRNA genes related to the Wnt signaling pathway (*WIF1*, *WNT2B*), solute carrier (SLC) gene family (*SLC9A3*, *SLC16A4*), and circadian rhythm (*KLF9*, *SIX3*).

**Figure 2.**
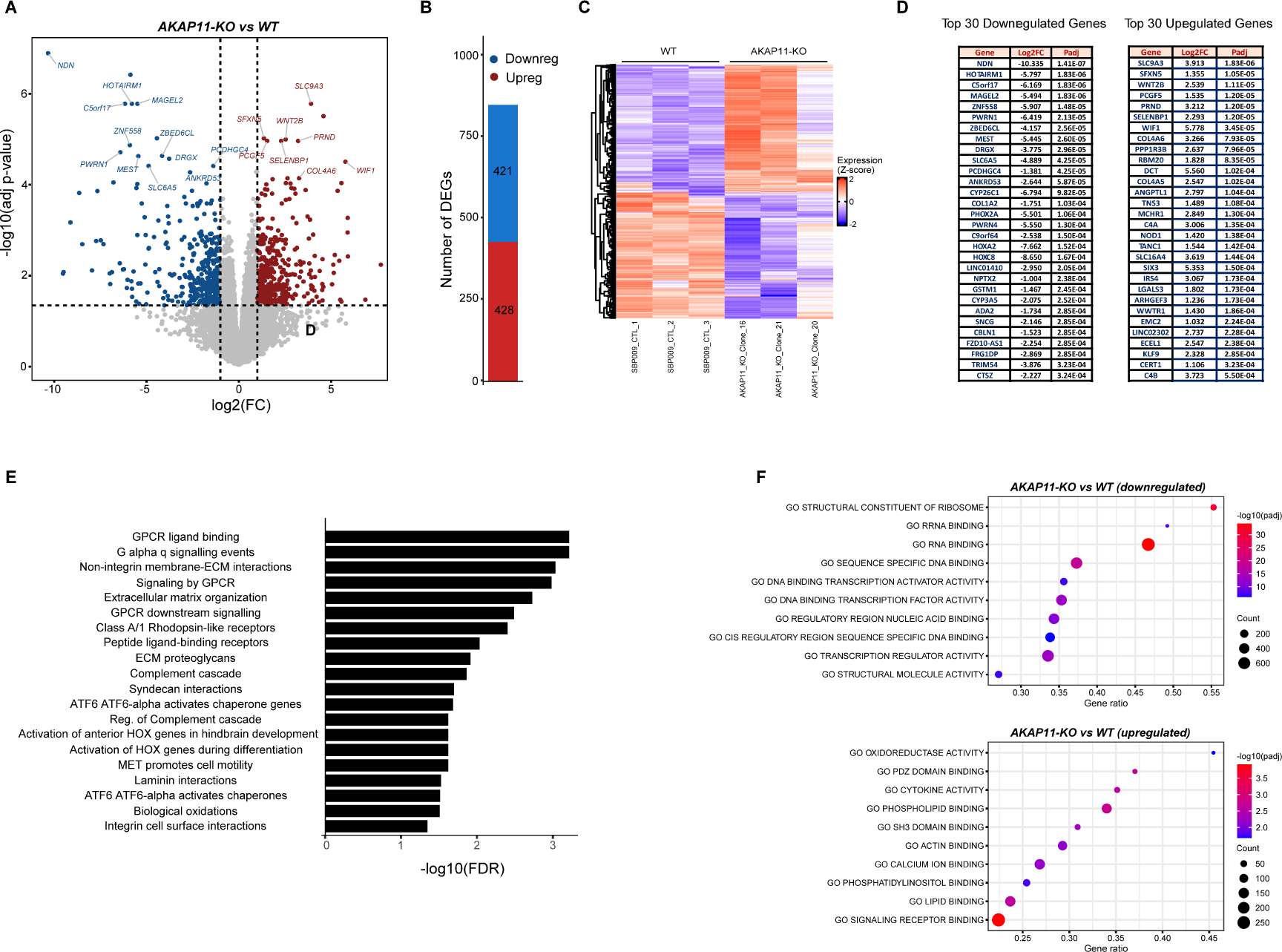
Identification of DEGs in heterozygous *AKAP11*-KO vs. WT and the pathways/GO terms affected. **A**) Volcano plot of differentially expressed genes (p-adj < 0.05 and |log2FC| > 1). **B**) Number of up- and down-regulated DEGs. **C**) Heatmap showing hierarchical clustering of differentially expressed genes (rows) in heterozygous *AKAP11*- KO and WT samples. The expression levels across samples were standardized by Z-Score method. **D**) List of the top 30 downregulated and upregulated DEGs in heterozygous *AKAP11-*KO vs. WT. **E**) Over-representation analysis (ORA) of all DEGs, both up- and downregulated, based on Curated. Reactome database using ShinyGO0.77; FDR < 0.05. The background genes used were all the 23,590 genes that were considered expressed (see Methods). Shown here are the top 20 pathways. For the full list refer to Supplementary Figure 2A. **F**) GSEA showing enrichment of GO (molecular function) terms in upregulated and downregulated genes from heterozygous *AKAP11-*KO vs. WT comparison; Padj<0.05. Shown here are the top 10 pathways. For the full list refer to Supplementary Table 2. The size of the circles represents gene count (count of core enrichment genes); the colors represent the strength of -log (adjusted p-value), with red showing the most significant adjusted p-value, and gene ratio represents (count of core enrichment genes) / (count of pathway genes).

We performed over-representation analysis (ORA) using Curated.Reactome^52,53^ pathway database to gain a general overview of the main biological pathways that were dysregulated using the obtained DEGs of the heterozygous *AKAP11-*KO vs. WT. We found that the DEGs were highly enriched in GPCR binding and signaling events as well as extracellular matrix (ECM) interactions and focal adhesion-related terms (Figure 2D). The genes that were enriched in the Curated.Reactome term “GPCR ligand binding” included Wnt/Frizzled receptors (*WNT8B*, *WNT2B*, *FZD10*), metabotropic glutamate receptors (GRM3), and Rhodopsin-like receptors (*UTS21*, *CXCL122*, *CCKBR2*, *PTGFR2*, *MCHR12*, *NTS2*, *HTR1B2*, *F2RL22*, *OXGR12*, *BDKRB22*, *GPR372*, *SSTR12*, *CRH2*), among others (Supplementary Figure 2A). GPCRs are the largest family of metabotropic receptors which play a pivotal role in neural communication by regulating synaptic transmission at both pre- and post-synaptic stages^54^.

To further extract biological meanings from the gene expression changes resulting from heterozygous *AKAP11-*KO and have a better understanding of their direction of change (up- or downregulated), we performed gene set enrichment analysis (GSEA^55,56^), which ranks the entire gene expression dataset based on t-statistic of each gene’s differential expression with respect to the two phenotypes of heterozygous *AKAP11*-KO vs. WT (Figure2F). The entire ranked list was used to evaluate how the genes from the Gene Ontology (GO), molecular function^57,58^ are overrepresented at the top or bottom of the ranked list of genes^59^. We showed that, overall, the main downregulated terms were related to ribosome function and protein synthesis as well as transcription regulation, particularly through changes in transcription factor binding activity (Figure 2F, Supplementary Table 2). Gene within the terms related to ribosome function and protein synthesis were mainly ribosomal protein genes and mitochondrial ribosomal protein genes. Furthermore, the transcription factors with differentially decreased expression in heterozygous *AKAP11*-KO were primarily from the family of HOX, Zinc Finger (ZNF), Iroquois homeobox (Irx), Activating transcription factor (ATF), and Paired-box (PAX) (Supplementary Table 2). On the other hand, the top-upregulated GO molecular functions in heterozygous *AKAP11*-KO were involved in diverse molecular functions including signaling receptor binding, phospholipid binding, postsynaptic Density-95 Discs-large Zona occludens 1 (PDZ)-domain binding, lipid binding, as well as calcium ion- and actin-binding (Figure 2F, Supplementary Table 2). Several of these functions, such as actin-binding, calcium ion binding, SRC Homology 3 (SH3) domain binding, lipid binding, and PDZ domain binding, include genes involved in cell adhesion, cell motility, or cytoskeletal integrity. Overall, the upregulated terms highlighted here are integral to signaling processes, receptor interactions, and cytoskeletal dynamics and are essential for the proper functioning of neurons.

### Genome-wide DMR profiling of heterozygous *AKAP11*-KO neurons compared to WT counterparts

We next performed WGBS using the three heterozygous *AKAP11*-KO clones and the three WT counterpart replicates and analyzed the genome-wide CpG DNA methylation modifications resulting from heterozygous *AKAP11*-KO. We detected the genome-wide DMRs (methylation values and p-values were averaged within each DMR), annotated them using Homer^47^ (DMRs = Peaks), and identified the genes associated with them (referred to as DMR-associated genes). We found 813 total DMRs and 705 total DMR-associated genes in heterozygous *AKAP11*-KO compared to WT—some genes have multiple DMRs of both directions of changes (Supplementary Table 3). Our data indicates that, overall, the methylation differences within the DMRs of heterozygous *AKAP11*-KO vs. WT are relatively small in magnitude, which is consistent with the notion of DNA methylation being a more stable epigenetic mark, that tends to remain more intact once it is established, especially in differentiated cells^60^. Among these DNA methylation modifications in DMRs, we note that heterozygous knockout of *AKAP11* leads to more hypermethylated DMRs than hypomethylated ones (638 vs. 175, respectively; Figure 3A). The hierarchical clustering and heatmap of DMRs, based on the mean methylation level β value in 3 heterozygous *AKAP11*-KO and 3 WT replicates, distinguishes the large fraction of DMRs with elevated methylation status from the smaller group with decreased methylation status in heterozygous *AKAP11*-KO vs. WT. Labeled on the heatmap are the top 10 hyper- and hypo-methylated DMR-associated genes. These include genes with various roles in brain development, neural regulation, and function, including transcription factors (*OTX1*, *OTX2*, *NR2E1*, *IRX2*, and *PAX7*), neurotransmission and signaling (*ENPP2*, *GNG11*, *GREB1L*, *NAALADL2*), and other regulatory molecules and structural components (Figure 3B, Supplementary Figure 3A). The majority of DMRs resulting from heterozygous *AKAP11*-KO fall within the non-coding intergenic and intronic regions, suggesting their potential involvement in non-promoter transcription regulation (Figure 3C, Supplementary Figure 3B).

**Figure 3.**
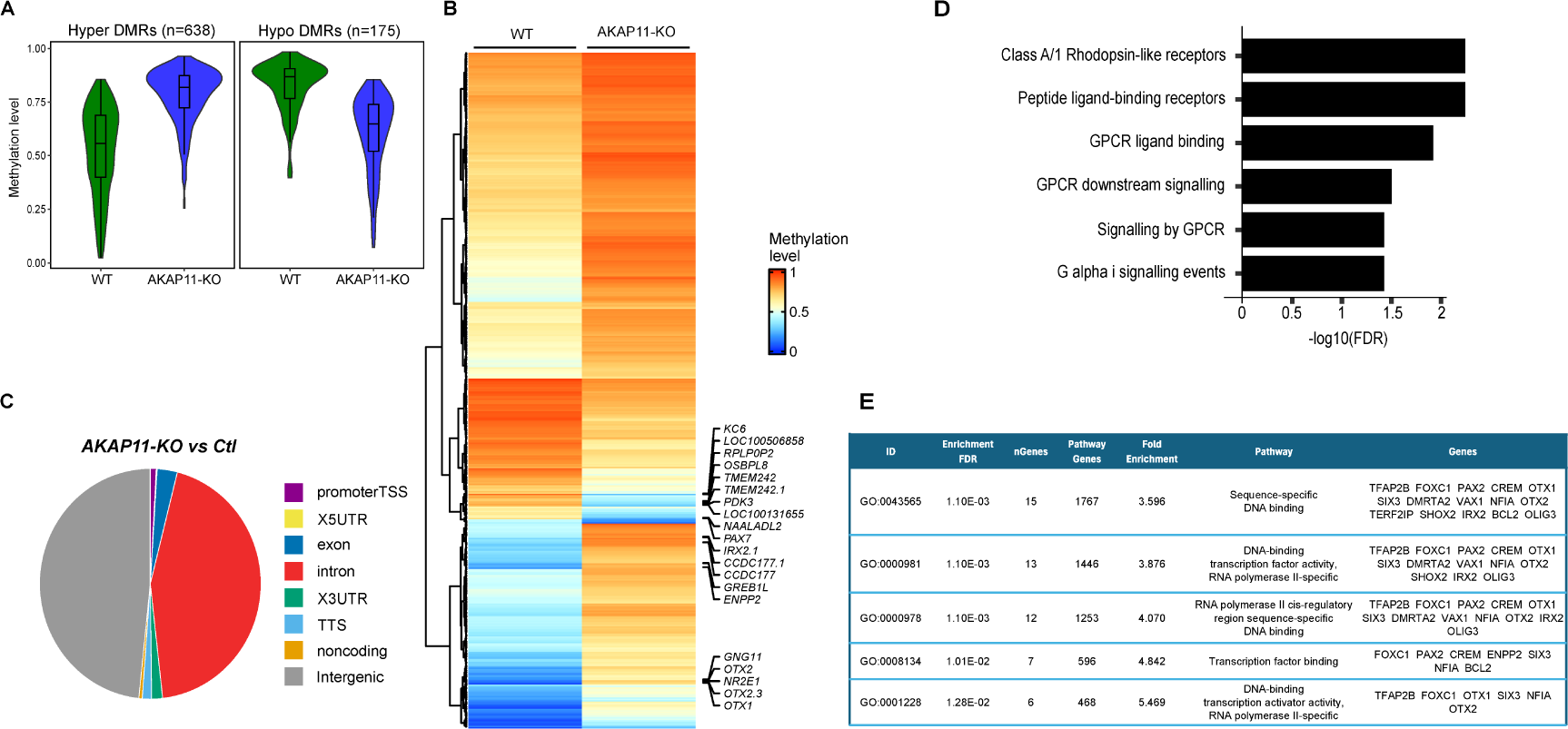
DMRs profiling of heterozygous *AKAP11*-KO compared to WT human iPSC-derived neurons. **A**) Violin plot for the overall distribution of methylation levels of hyper- and hypo-DMRs in heterozygous *AKAP11*-KO (colored blue) and WT (colored green) conditions (see Supplementary Table 3 for complete the list of DMRs). Methylation levels in WT and KO were obtained from the same hyper- or hypo-DMRs. Hypermethylation= positive methylation difference, Hypomethylation = negative methylation difference. **B**) Heatmap of patterns of methylation constructed using unsupervised hierarchical clustering of all DMRs identified (rows); columns represent the average of 3 WT reps and 3 KO clones. Orange and blue colors show the highest and lowest methylation levels, respectively. **C**) Pie chart of genomic distribution of 813 DMRs across the genome, illustrating the proportions of genomic features. **D**) Enrichment analysis (ORA) of all genome-wide DMRs, regardless of their direction of change, using Curated.Reactome databases. Generated using ShinyGO0.77; FDR < 0.05; top 20 pathways. **E**) Enrichment analysis (ORA) of genome-wide DMRs that are associated with gene expression changes (regardless of their direction; P-adj < 0.05 & |log2FC| > 0.25), using GO molecular function database. Generated using ShinyGO0.77; FDR < 0.05; top 5 pathways (redundancy removed). For the full top 20 pathways refer to Supplementary Figure 3D.

Subsequently, in search for over-represented pathways among all DMR-associated genes in heterozygous *AKAP11*-KO vs. WT, regardless of their direction of change, we performed ORA using Curated.Reactome database. We detected that the DMRs are primarily enriched in terms such as Class A/1 Rhodopsin−like receptors, peptide ligand-binding receptors, GPCR ligand binding, and GPCR downstream signaling (Figure 3D, Supplementary Figure 3C). Interestingly, the dysregulation of the GPCR-related signaling pathway in heterozygous *AKAP11*-KO obtained from the ORA of DMRs is in line with our findings from the ORA of DEGs. Lastly, using all these DMR-associated genes that were correlated with differential gene expression changes (59 genes; DEGs: P-adj < 0.05 & |log2FC| > 0.25), we performed GO molecular function enrichment analysis to uncover biological meaningful roles of specifically those DMR-associated genes that are linked with gene expression changes. The top 10 enriched GO terms highlight DNA-binding transcription factor activity (Figure 3E, Supplementary Figure 3D, Supplementary Table 4). While global DMR-associated genes were enriched in GPCR binding and signaling terms, those that were significantly linked with the gene expression modifications highlighted the enrichment of DNA binding transcription factor activity.

Overall, we show that in heterozygous *AKAP11*-KO compared to WT, the methylation difference values of the DMRs are quite subtle in magnitude, there is a higher number of hypermethylated DMRs than hypomethylated ones, and the DMRs are mainly residing within the intergenic and intronic regions. We also show that the global DMR-associated genes are enriched in pathways related to GPCRs’ function and binding, while those that are specifically associated with differential gene expression modifications primarily function as DNA-binding transcription factors.

### Enhancer activity dysregulations following heterozygous *AKAP11*-KO play a key role in the aberrant expression of neighboring genes

Based on our results described above that highlighted aberrant expression of genes related to transcription factor activity, as well as predominant distribution of DMRs in intergenic and intronic regions, we inferred potential dysregulation of transcription factor binding activity at cis-regulatory regions—mainly enhancers that play a role in aberrant gene regulatory networks. To test this hypothesis, we performed ChIP-seq for H3K27ac, marker of active enhancers^61-63^, and investigated the modifications in heterozygous *AKAP11*-KO. Our data show that H3K27ac peaks with decreased intensities in heterozygous *AKAP11*-KO are preferentially located in intergenic regions while those with increased intensity present a more similar distribution to the set of all consensus peaks, with a relatively higher distribution in intronic regions (Figure 4A). Overall, in the intergenic regions, the number of H3K27ac differential peaks with reduced intensities in heterozygous *AKAP11*-KO was almost twice the number of differential peaks with increased intensities (6,763 down and 3,612 up; differential peaks at adj-P < 0.05 & |log2FC| > 1; Figure 4B). In the intronic regions, however, the difference between the number of increased and decreased differential peaks is less pronounced, yet more differential peaks show increased intensities in these regions (5719 down and 6146 up; Figure 4B).

**Figure 4.**
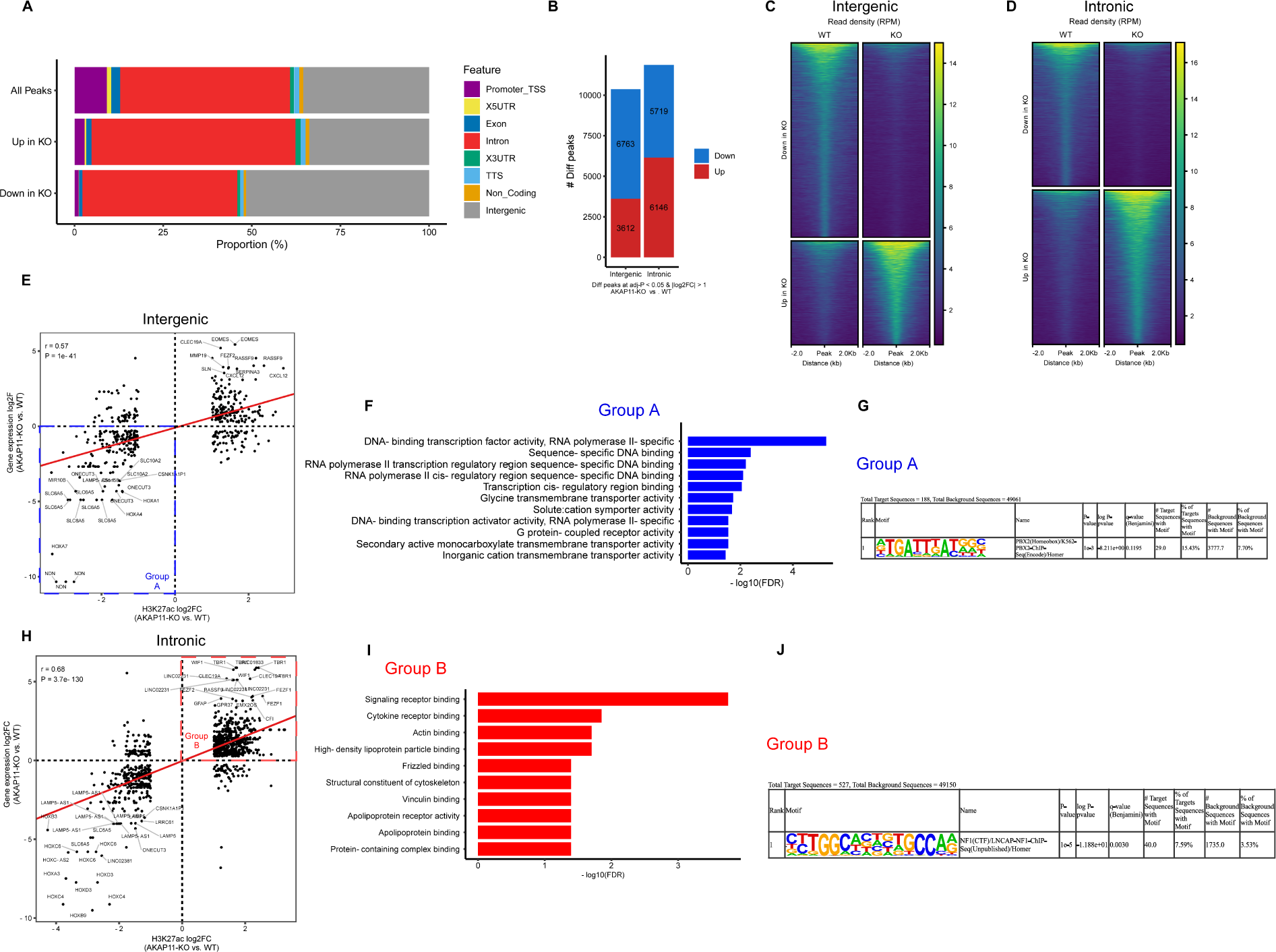
Intergenic and Intronic differential H3K27ac peaks profiling and investigating the association between H3K27ac level modifications and gene expression changes of target genes. **A)** Distribution of genomic compartments overlapping different subsets of H3K27ac peaks categorized by differential binding status; All peaks= all consensus peaks; <u>Up in KO</u>: differential H3K27ac peaks with increased intensities in heterozygous *AKAP11*-KO vs. WT; <u>Down in KO</u>: differential H3K27ac peaks with reduced intensities in heterozygous *AKAP11*-KO vs. WT. **B)** Number of differential H3K27ac peaks with increased (up) and decreased (down) intensities in heterozygous *AKAP11*-KO vs. WT; adj-P < 0.05 & |log2FC| > 1. **C, D)** Heatmap of intergenic and intronic differential H3K27ac signal enrichment (heterozygous *AKAP11*-KO and WT) centered at the center of peaks, ±2 kb flanking region. Rank ordered by intensity of H3K27ac peaks based on reads per million mapped reads (RPM). For the heat map of individual samples and average read density profiles refer to Supplementary Figure 4A-D. **E)** Correlation plot of differential gene expression fold changes (adj-P < 0.05 & |log2FC| > 0.25) and differential intergenic H3K27ac fold changes (adj-P < 0.05 & | log2FC| > 1), using a 100 kb distance cutoff. r = 0.57, p = 1e−41. **F)** GO-Molecular Function ORA of reduced intergenic H3K27ac peak intensities associated with decreased gene expression of target genes in heterozygous *AKAP11*-KO vs. WT (intergenic H3K27ac down, expression down: group A. Generated using ShinyGO-0.77; FDR < 0.05; top 20 pathways demonstrated (redundant terms removed). For the full top 20 list refer to Supplementary Figure 4E. **G)** Top transcription factor binding sites query result for group A, using HOMER motif analysis (Known Motif Enrichment). Total Target Sequences = 188, Total Background Sequences = 49061. p= 1e-3. For the full table refer to Supplementary Table 6. **H)** Correlation plot of differential gene expression fold changes (adj-P < 0.05 & |log2FC| > 0.25) and differential intronic H3K27ac fold changes (adj-P < 0.05 & |log2FC| > 1), using a 100 kb distance cutoff. r = 0.68, p = 3.7e−130. **I)** GO-Molecular Function ORA of increased intronic H3K27ac peak intensities associated with increased gene expression of target genes in heterozygous *AKAP11*-KO vs. WT (intronic H3K27ac up, expression up: group B). Generated using ShinyGO-0.77; FDR < 0.05; top 20 pathways demonstrated. **J)** Top transcription factor binding site query result for group B, using HOMER motif analysis (Known Motif Enrichment). Total Target Sequences = 527, Total Background Sequences = 49150. p=1e-5. For the full table refer to Supplementary Table 6.

Using heatmaps, we next separately profiled the intergenic and intronic H3K27ac enrichment patterns in heterozygous *AKAP11*-KO and WT centered at the center of the peaks, ±2 kb flanking region (signal from KO clones and WT replicates were averaged using WiggleTools^64^). Data were normalized so that each value represents the read count per base pair per 10 million reads. The heatmaps not only indicate a clear distinction between the enrichment of H3K27ac signals between heterozygous *AKAP11*-KO and WT in intergenic and intronic regions but also confirm the higher enrichment of intergenic H3K27ac peaks with differentially decreased intensities (down) and intronic peaks with differentially increased intensities (up) in heterozygous *AKAP11*-KO (Figure 4C and D; Supplementary Figure 4A-D). As H3K27ac marks active enhancers and active chromatin^61-63^, and is indicative of enhancer activity^65^, we utilized this chromatin signature to detect active enhancers and predict their target genes by linking the H3K27ac peaks with the nearest TSS of known genes. Acknowledging that the nearest assigned gene to the enhancer might not necessarily be the true target of that enhancer and that these enhancers might not be the main determinant of the gene expression fate of the target genes, we investigated the correlation between differential H3K27ac peaks changes (P < 0.05 & |log2FC| > 1) and differential gene expression modifications (adj-P < 0.05 & |log2FC| > 0.25) in intergenic and intronic regions. We found a significant positive correlation in both cases (intergenic: r = 0.57, p = 1e−41; intronic: r = 0.68, p = 3.7e−130; Figure 4E and H, Supplementary Table 5). To determine which set of target genes are mainly affected by the enhancers in our system, and as we already unraveled that enhancers with differentially decreased strength in heterozygous *AKAP11*-KO were preferentially mapped to intergenic regions and those with differentially increased intensities were more abundant in intronic regions, we narrowed down our focus to two groups of enhancers: 1. <u>Group A</u> (intergenic differential H3K27ac down, differential expression down; 105 genes): Intergenic enhancers with decreased differential activity in heterozygous *AKAP11*-KO whose target genes show a negative differential gene expression fold change (Figure 4E), 2. <u>Group B</u> (intronic differential H3K27ac up, differential expression up; 242 genes): Intronic enhancers with increased differential activity in heterozygous *AKAP11*-KO whose target genes show a positive differential gene expression fold change (Figure 4H). To identify the functional properties of target genes within these two groups, we performed ORA using the GO molecular function database. For group A, the top GO terms were primarily related to DNA-binding transcription factor activity, highlighting that reduced intergenic enhancer activity upon heterozygous knockout of *AKAP11*, at least partially, accounts for the downregulation of genes involved in regulatory networks that play key roles as transcription factors or co-factors, as previously observed in our gene expression and DMR analysis (Figure 4F, Supplementary Figure 4E). The top GO terms in this group also include glycine transmembrane transporter activity (*SLC32A1* and *SLC6A5*), solute:cation symporter activity (*SLC32A1*, *SLC5A7*, *SLC10A2*, *SLC6A5*), and G protein-coupled receptor activity (*ADGRF5*, *ADCYAP1R1*, *HRH3*, *FZD10*, *CXCR4*, *SSTR1*, *GPRC5C*; Figure 4F, Supplementary Figure 4E). For group B the top GO terms involve signaling receptor binding, cytokine receptor binding, actin binding, high-density lipoprotein particle binding, frizzled binding, and structural constituent of cytoskeleton, among others (Figure 4F, Supplementary Figure 4F).

In order to detect the transcription factor binding motifs involved in the down- and upregulation of the genes enriched in GO terms (Group A and B), we performed transcription factor binding site (TFBS) motif analysis using HOMER^47^. Scanning the H3K27ac differential peaks (adj-P < 0.05 & |log2FC| > 1) that were associated with differential gene expression (adj-P < 0.05 & |log2FC| > 0.25) in the concordant direction for known TFBS motifs, we detected the most significant enrichment to be PBX2/Homer in group A (p=1e-3, 15.43% of target peaks, 7.70% of background) and NF1/Homer in group B (p=1e-5, 7.59% of target peaks, 3.53% of background).

### Intergenic and intronic differential enhancer activity is negatively correlated with methylation difference within DMRs in the respective enhancer regions in heterozygous *AKAP11*-KO vs. WT

To comprehensively explore the interplay between both intergenic and intronic enhancer activity, marked by H3K27ac, and alterations in DNA methylation levels at those regions, following the heterozygous knockout of *AKAP11*, correlation plots were constructed separately for intergenic and intronic regions, depicting the relationship between the differential H3K27ac log2FC (heterozygous *AKAP11*-KO vs. WT, adj-P < 0.05 & |log2FC| > 1) and methylation differences within the DMRs. Notably, we identified significant negative correlations between the subtle methylation differences in DMRs and differential H3K27ac changes in both intergenic and intronic regions (r = −0.52, P = 0.00022 and r = −0.37, P = 0.019, respectively; Figure 5A and B, Supplementary Table 7). This observation aligns with the established pattern where activated enhancers typically display a distinct reduction in DNA methylation (5mC), a phenomenon observed across various tissues and cell types^68-71^.

**Figure 5.**
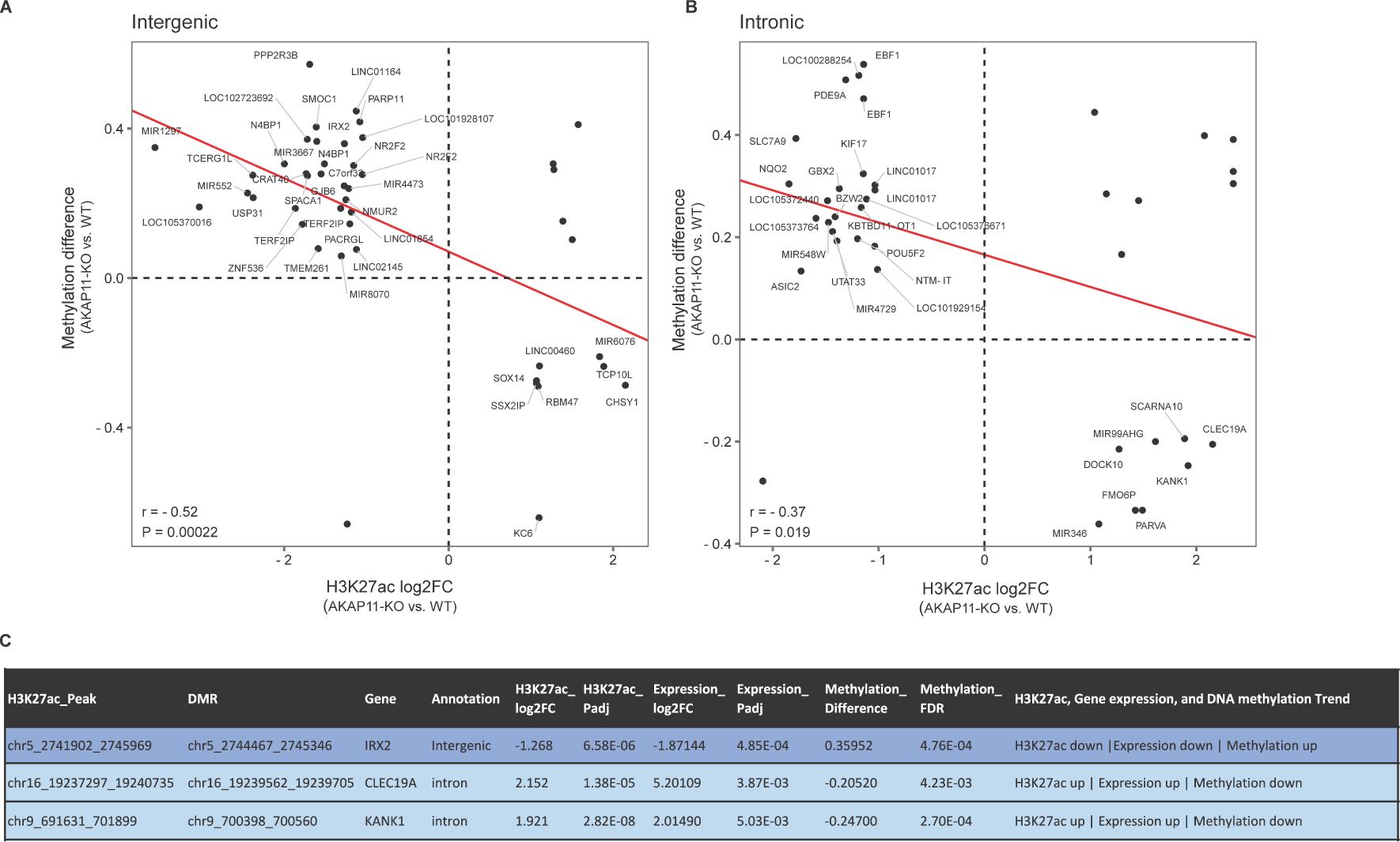
DMR status in enhancers with concordant changes in the gene expression of target genes (using H3K27ac, DNA methylation, and gene expression datasets). **A, B**) Correlation plots of differential H3K27ac log2FC (adj-P < 0.05 & |log2FC| > 1) and methylation difference within DMRs for *AKAP11*-KO vs. WT in intergenic (r = −0.52, P = 0.00022) and intronic regions (r = −0.37, P = 0.019). **C**) A small subset of enhancers with differential activity with concordant differentially expressed target genes in *AKAP11*-KO showing significant negative correlation with DMRs at those enhancers. H3K27ac and Gene expression cutoffs: adj-P < 0.05 & |log2FC| > 1; DMR: callDMR at default parameters with a minimum length of 50 base pairs and 3 CpG sites.

Employing a multi-omics approach, we leveraged data on DEGs (cutoff: adj-P < 0.05 & |log2FC| > 1), DMRs (called as previously described in methods), and differential H3K27ac (cutoff: adj-P < 0.05 & |log2FC| > 1) to pinpoint the most profoundly impacted intergenic and intronic regulatory regions influenced by heterozygous LoF mutations in *AKAP11*. Given the overall negative correlation between DMR methylation differences and differential H3K27ac fold changes in our system, along with previous findings suggesting that transcription factor binding at methylated regions is linked with a reduction in DNA methylation, at least in a subtype of enhancers^68,72^, we focused on differential enhancers exhibiting an inverse relationship with DMRs and concurrent differential expression of target genes. This analysis revealed that, in heterozygous *AKAP11*-KO, aberrantly methylated regions in intronic and intergenic enhancers with differential activity and an inverse direction of change significantly affected 3 regions that were linked to concordant gene expression alterations in the neighboring genes. These enhancers, tied with chromatin state changes and gene expression regulation of target genes, were mapped to Iroquois Homeobox 2 (*IRX2)*, C-type lectin domain containing 19A (*CLEC19A)*, and KN Motif And Ankyrin Repeat Domains 1 (*KANK1;* Figure 5E).

## Discussion

Given the complex polygenic nature of BD and SCZ, as well as our limited understanding of their origins and etiology, it is important to study novel risk genes. Genetic studies identified AKAP11 LoF as a candidate predisposing gene for both diseases. A study looking at AKAP11 mutant mice found that their EEG patterns resembled those observed in individuals with SCZ^31^. In another study, proteomic profiling of synapses found shared molecular pathway modifications between patients with SCZ, BD, and *AKAP11*-mutant mouse models--mainly pathways related to ribosomes, mitochondrial respiration, and vesicle trafficking pathways^34^. Our study, for the first time, uncovers the transcriptomic and epigenomic consequences of heterozygous LoF mutations in *AKAP11*, a novel rare-variant large-effect risk gene for BD and SCZ, in the context of human iPSC-derived neurons.

AKAP11 can modulate gene expression by influencing the activity of transcription factors through its role in PKA signaling. PKA has been implicated in a multitude of signaling processes and has an extensive array of downstream targets and processes^76^ such as phosphorylating transcription factors (e.g. CREB), neurotransmitter receptors (e.g., AMPA receptors), cytoskeleton-associated proteins, ion channels, and elements of other signaling cascades^77,78^. Interestingly, our transcriptomic analysis found dysregulation of genes involved in many of these functions in our heterozygous *AKAP11*-KO neuronal model. Heterozygous LoF mutations in *AKAP11* may lead to reduced PKA activity^79^ and as PKA regulates transcription factors, directly, or indirectly through binding to other kinases and substrates, its reduced activity could potentially lead to impaired activation of certain transcription factors and thus reduced expression of numerous genes that can themselves function as transcription factors or co-factors, as highlighted by our results. Particularly, in the case of CREB, its downstream target genes are involved in various processes, including metabolism (e.g., cytochrome c, *PEPCK*, and *aminolevulinate synthase*), transcription (e.g., *ATF-3*, *STAT3*, and *c-fos*), cell survival (e.g., *bcl-2*, *cyclin D1*, and c*yclin A*), and growth factors (e.g., *BDNF* and *FGF6*). CREB alone can regulate more than 4,000 genes^24,80^, many of which are themselves transcription factors (e.g., *NF-IL6*, *ZNF268*, and *NR4A2*)^81^. Considering that phosphorylation of transcription factors can positively or negatively affect their activity^82^, the wide range of genes with significantly up-or down-regulated profiles with their diverse roles in neurons could be a direct or indirect result of aberrant PKA-dependant phosphorylation of transcription factors.

Our findings also suggest that the downregulation of genes involved in DNA-binding transcription factor activity can be partly explained by the reduced activity of their intergenic enhancers, which indicates the involvement of histone acetyltransferases (HATs) and/or HDACs and chromatin accessibility changes at such regulatory regions. The acetylation and deacetylation of histones, facilitated by HATs and HDACs, respectively, play a role in regulating gene transcription^73,83,84^. It has been shown that the AKAP11-involved cAMP/PKA signaling inhibits gene transcription by phosphorylating HDACs, such as HDAC5, and preventing their export from the nucleus^25^. In addition, AKAP11-PP1 complex can affect HDAC6 stability^85^. Moreover, AKAP11-PKA-regulated GSK3B, one of the molecular targets of lithium, which is the most common medication to prevent recurrences of both manic and depressive episodes in BD patients^18-21^, has been suggested to be involved in the regulation of HDAC6 activity through phosphorylation events^26^. On the other hand, CREB binding protein (CREBBP), which is the transcriptional coactivator of CREB and is also phosphorylated by PKA, can function as an intrinsic HAT^86,87^. The DMR-associated genes that overlap with gene expression modifications in heterozygous *AKAP11*-KO also refer to transcription factor activity as the top GO term, suggesting that DMRs play partial roles in such gene expression dysregulations in heterozygous *AKAP11*-KO.

Consistent with previous research providing evidence of downregulated mitochondrial ribosomal subunit genes in the brain and blood samples of SCZ patients^88^ as well as decreased synaptic mitochondrial content and ribosomal proteins in patients with BD, SCZ, and *AKAP11*-mutant mice^34^, we detected significant downregulation of ribosomal protein genes and mitochondrial ribosomal protein genes along with other regulators involved in protein synthesis in our model. One possible explanation for the decreased expression of ribosomal proteins following heterozygous KO of *AKAP11* could be the inhibition of mTORC1 upon modifications in the upstream signals including growth factors, stress, or energy status^89^. Mitochondrial ribosomal proteins, particularly, are involved in mitochondrial oxidative phosphorylation (OXPHOS)^90^. It has been reported that the absence of *AKAP11* interrupts mitochondrial respiration by blocking the PKA activity^79^. Furthermore, synaptic mitochondria content supplies energy for local translation in the process of synaptic plasticity^91,92^. As ribosomal proteins are predominantly located in neurites, downregulation of ribosomal protein genes may lead to reduced synthesis of synaptic proteins^93^ as a part of a plasticity mechanism where neurons adjust their excitability in response to network activity^93^, an event that occurs in neuropsychiatric disorders^94,95^.

We found that heterozygous knockout of *AKAP11 led to* upregulation of pathways such as actin-binding, calcium ion binding, SH3 domain binding, lipid binding, and PDZ domain binding, which include genes involved in cell adhesion, motility, ECM interactions, or cytoskeletal integrity--pathways that have been previously suggested to be associated with lithium response^96,97^. Moreover, we found that the upregulated genes that were linked to increased intronic enhancer activity were primarily enriched in similar pathways suggesting that intronic enhancer activity through HATs and/or HDACs, plays a role, at least in part, in such dysregulations. It is known that generally, IQ domain GTPase-activating proteins (IQGAPs) play a role in remodeling the cytoskeleton, cell-cell adhesion, and regulating cell growth^22,98,99^. Functional studies have reported that *AKAP11* and the cytoskeletal scaffolding proteins IQGAP1/2 interactions affect cell motility^100^ and actin cytoskeleton^22^. In addition, *AKAP11* inhibits GSK3B, and GSK3, through its substrates, can regulate actin, tubulin, and the cytoskeleton structure^100^. All these findings support the involvement of *AKAP11* in the regulation of cell motility and actin cytoskeleton possibly through interacting with GSK3 and IQGAPs, which may account for the transcriptomic aberrations seen in the knockout of *AKAP11*. ECM dysregulations and decreased actin-related gene expression have been reported in the post-mortem cortex studies of SCZ patients^101,102^. Impaired adhesion is closely connected to both cell migration and axon guidance, pathways that exhibit abnormalities in genes associated with the risk of SCZ^103-105^. Additionally, there is a BD and SCZ-specific association observed in pathways related to the dynamic regulation and restructuring of cytoskeletal actin^106,107^, cytoskeletal proteins^108^, and cellular communication with ECM^109^. Lastly, our GSEA results highlight cytokine activity upregulation in heterozygous *AKAP11*-KO and the follow-up analysis of upregulated genes associated with significantly increased intronic enhancer activity suggests that enhancer activity alterations and potential involvement of histone HATs and/or HDACs partially explain the dysregulation of transcription of the genes involved in cytokine receptor binding. Cytokine activity in the brain can affect mood, memory, cognition, and sleep and modulate neuroendocrine stress responses^110^. Notably, literature evidence highlights a potential causative role for cytokines in SCZ development^111^ as well as a partial overlap in the increased blood cytokine levels in patients with BD with psychotic symptoms and first-episode SCZ^112^. Most importantly, dysregulated blood levels of cytokine network elements have been detected in BD, SCZ, and major depressive disorder ^113-119^.

Non-promoter regulatory elements are rich in factors that contribute to neuropsychiatric heritability^120-122^. It has been demonstrated that the levels of DNA methylation, H3K27ac, and overall chromatin accessibility in these areas can have effects on disorders specific to certain regions of the brain^120-122^. Moreover, numerous genetic risk loci associated with SCZ and BD have been previously reported to be within non-coding regions, including introns, intergenic regions, and noncoding RNA genes, indicating the pivotal involvement of gene regulation in the disease’s development^73-75^. Most importantly, BD and SCZ brains have previously demonstrated dysregulation of regulatory sequences, particularly modifications that affect the neuronal H3K27ac levels^123^. Our findings from intergenic and intronic H3K27ac motif analysis show that, in heterozygous *AKAP11*-KO, the intergenic enhancers with reduced transcription factor binding activity that were linked to downregulation of target genes were mainly enriched in PBX2 binding motifs, suggesting a potential impairment of PBX2 transcription factor binding at intergenic enhancers and subsequent downregulation of target genes. *PBX2* (HOX12) encodes a widely expressed member of the TALE/PBX homeobox family and interacts with a specific group of HOX proteins, improving their capacities for DNA binding in terms of both affinities and specificities^124,125^. Genome-wide association studies have reported strong associations of PBX2 with the risk of SCZ and autism spectrum^126,127^. On the other hand, intronic enhancers with elevated transcription factor binding activity, linked with the upregulation of target genes, were significantly enriched in NF1 binding motifs, suggesting a potential role of NF1 binding at intronic enhancers in the upregulation of target genes. NF1 transcription factors can bind to specific DNA sequences with high affinity and are abundant in cerebellar granule neurons^128,129^. Their activity can be modulated post-transcriptionally, through phosphorylation^130-132^ or regulation of mRNA stability^130,133^. The DNA binding capability of a particular form of NF1 has been demonstrated to be dependent on ECM in mammary epithelial cells^134^.

Investigating the genome-wide DNA methylation modifications not only uncovered the overall genome-wide changes seen with the knockout of *AKAP11* but also, along with the transcriptomic data, showed that the DMRs were specifically linked to gene expression dysregulations. Such DMR-associated genes were primarily involved in transcription factor binding functions. Although DMR-associated genes showed enrichment of GPCR-related functions, further investigation found that DMRs did not have a considerable effect on transcription dysregulation of genes involved in GPCR-related functions. Our findings imply that a portion of GPCR-signaling-related gene expression modifications, notably *GPRC5C*, *FZD10*, and *WNT2B*, are associated with enhancer activity alterations, suggesting the potential involvement of HATs and/or HDACs. Altered G-protein-linked signaling system and its downstream targets have been traditionally implicated in the pathophysiology and pharmacology of BD, SCZ, and major depressive disorder by our group and others^135-140^. The observed overall negative correlation between intronic and intergenic methylation differences in DMRs and differential H3K27ac changes in those regions suggests that, for the most part, transcription factor binding at active enhancers is inversely related to DNA methylation status in those sites. Previous findings reported that transcription factor binding at methylated regions is associated with decreased DNA methylation^68,72^. Notably, beyond preventing aberrant activation of the genome, a recent study showed that DNA methylation plays a direct role in regulating transcription factor binding in a small subset of cell type-specific active enhancers^72^. We detected enhancers with differential activity, overlapping DMRs of the opposite direction of change, which were linked to concordant differential gene expression changes to be mapped to *IRX2, CLEC19A*, and *KANK1*. Generally, IRX transcription factors play roles in neuronal function and development^141^. IRX2, a transcriptional inhibitor, has been suggested to have a connection to social behaviors in animals ^142,143^. The expression of *IRX2* is known to be modulated by the three-dimensional structure of chromatin, which itself is controlled by the architectural protein known as CCCTC-binding factor (CTCF)^144-146^. Intergenic hypermethylation of *IRX2* noted by DMRs, along with a significant decrease in H3K27ac peaks in that region and its reduced gene expression level in heterozygous *AKAP11*-KO implies possible chromatin structure state changes and is in concordant with previous reports suggesting DNA methylation’s ability to compete with CTCF binding, particularly at CpGs in key regulatory regions^147,148^. KANK1 plays a pivotal role in cell motility and cytoskeleton formation through modulating actin polymerization^149,150^. Its activity has been shown to inhibit actin fiber formation, cell migration, RhoA activity, and neurite outgrowth^151^. Interestingly, the first large-scale meta-analysis of CNVs across SCZ, bipolar disease, depression, ADHD, and autism cohorts has detected enrichment of duplications in *KANK1* in all five groups^152^. Finally, CLEC19A, which belongs to the C-type lectin family, is highly expressed in brain tissue^152^. The *CLEC19A* gene is close to a locus that has been previously associated with attention deficit hyperactivity disorder (ADHD)^153^ and it has been suggested that there is common genetic etiology between ADHD, BD, and SCZ ^154^.

Overall, our investigations shed light on the impact of heterozygous *AKAP11*-KO on gene expression modifications, particularly genes playing roles as DNA-binding transcription factors, actin and cytoskeleton regulators, and cytokine receptors, which are at least partially dysregulated by enhancer activity alterations, marked by H3K27ac, as well as DNA methylation modifications detected within DMRs. We also highlighted the enrichment of GPCR binding and signaling within DEGs, genome-wide DMR-associated genes, and showed that a portion of those DEGs were linked to enhancer activity alterations. Further investigation is essential to explore the precise mechanism of action and functional significance of heterozygous *AKAP11*-KO within the pathways, roles, and epigenetic regulators linked to such modifications. This is imperative for advancing our comprehension of the roots and etiology of BD and SCZ as well as eventually finding the most appropriate therapeutic agents for patients.

## Supplementary Material

### Supplementary Figures

**Supplementary Figure 1.**
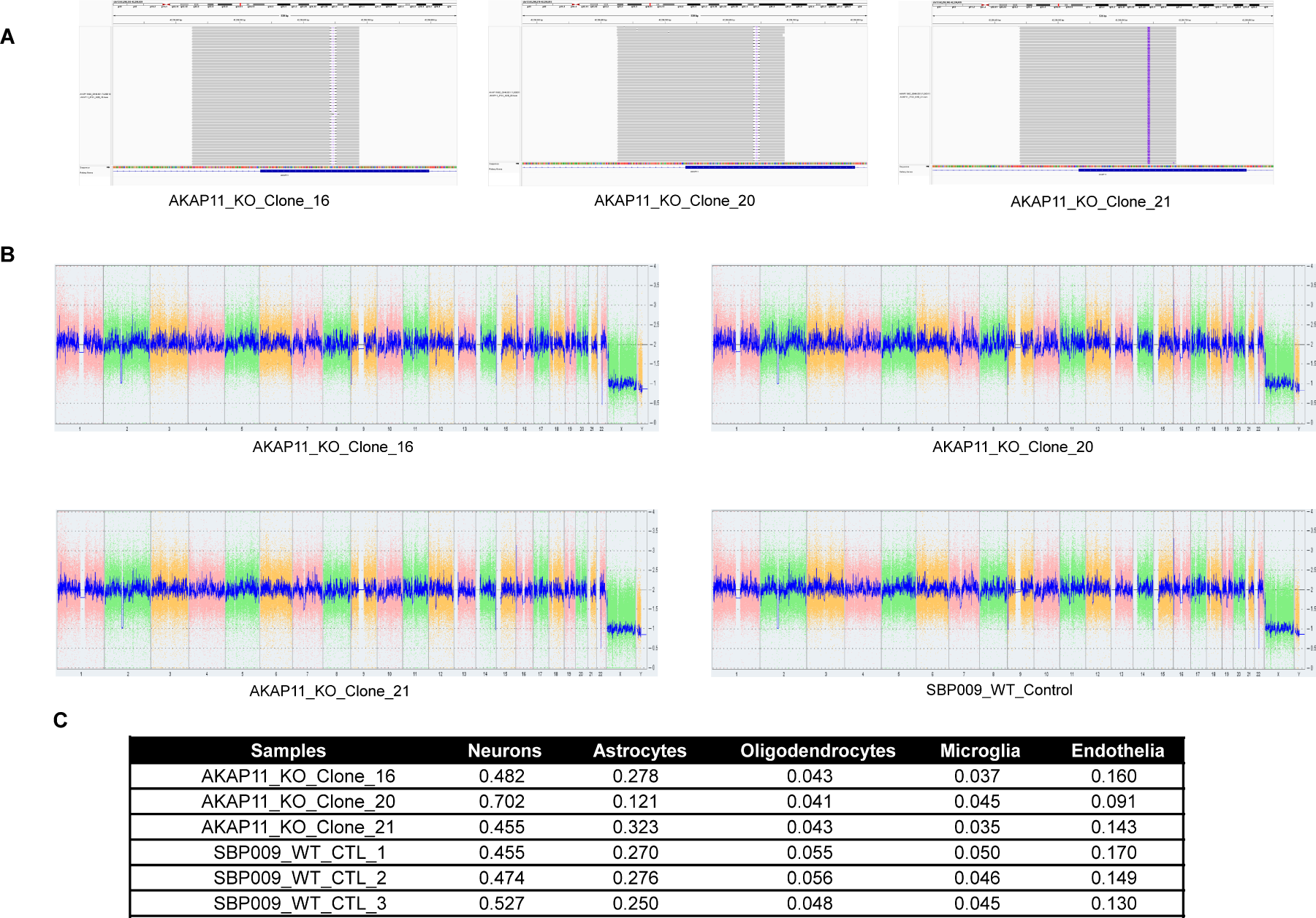
Validation of *AKAP11* heterozygous knockout in three isogenic clones, their chromosomal integrity, and abundance of neurons in the cultures. **A)** Miseq amplicon sequencing results showing heterozygous frameshift LoF mutations in iPSC *AKAP11*-KO clones. B) KaryoStat+ assay for digitally visualizing chromosomal aberrations. The size of structural aberration detectable is > 1 Mb for chromosomal gains and losses. The smooth signal plot is the smoothing of the log2 ratios which show the signal intensities of probes on the microarrayCN = 2. A value of 3 represents chromosomal gain (CN = 3). A value of 1 represents a chromosomal loss (CN = 1). C) Cell type deconvolution using bulk RNA-seq data and dtangle method^51^ from BrainDeconvShiny^155^, showing different cell types present in the cultures.

**Supplementary Figure 2.**
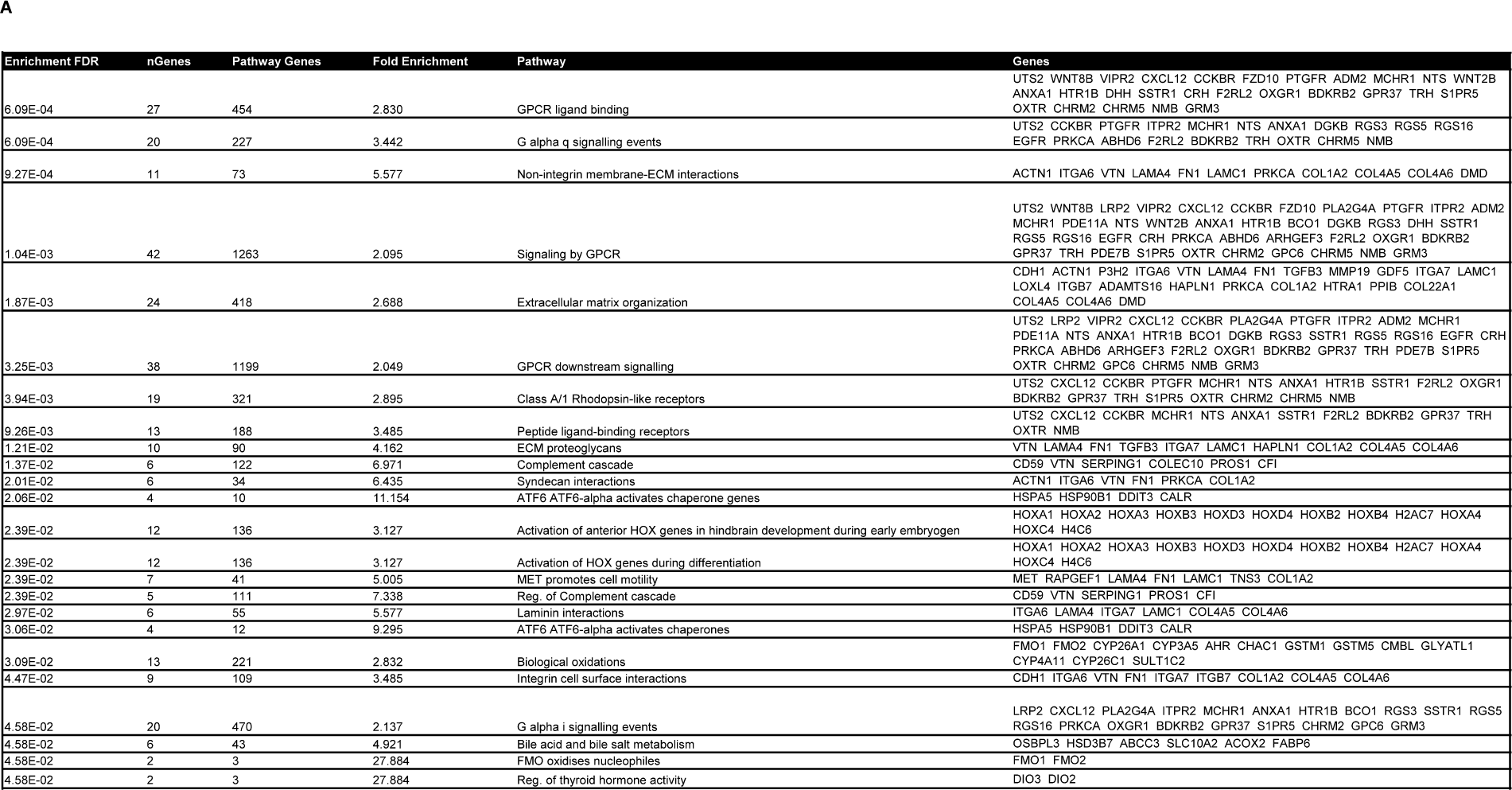
Results table of ORA using all DEGs. **A)** Table of overrepresented Reactome pathways resulting from ORA of DEGs (both up- and downregulated) of heterozygous *AKAP11*-KO vs. WT. Generated using ShinyGO 0.77; FDR < 0.05; top 20 pathways demonstrated.

**Supplementary Figure 3.**
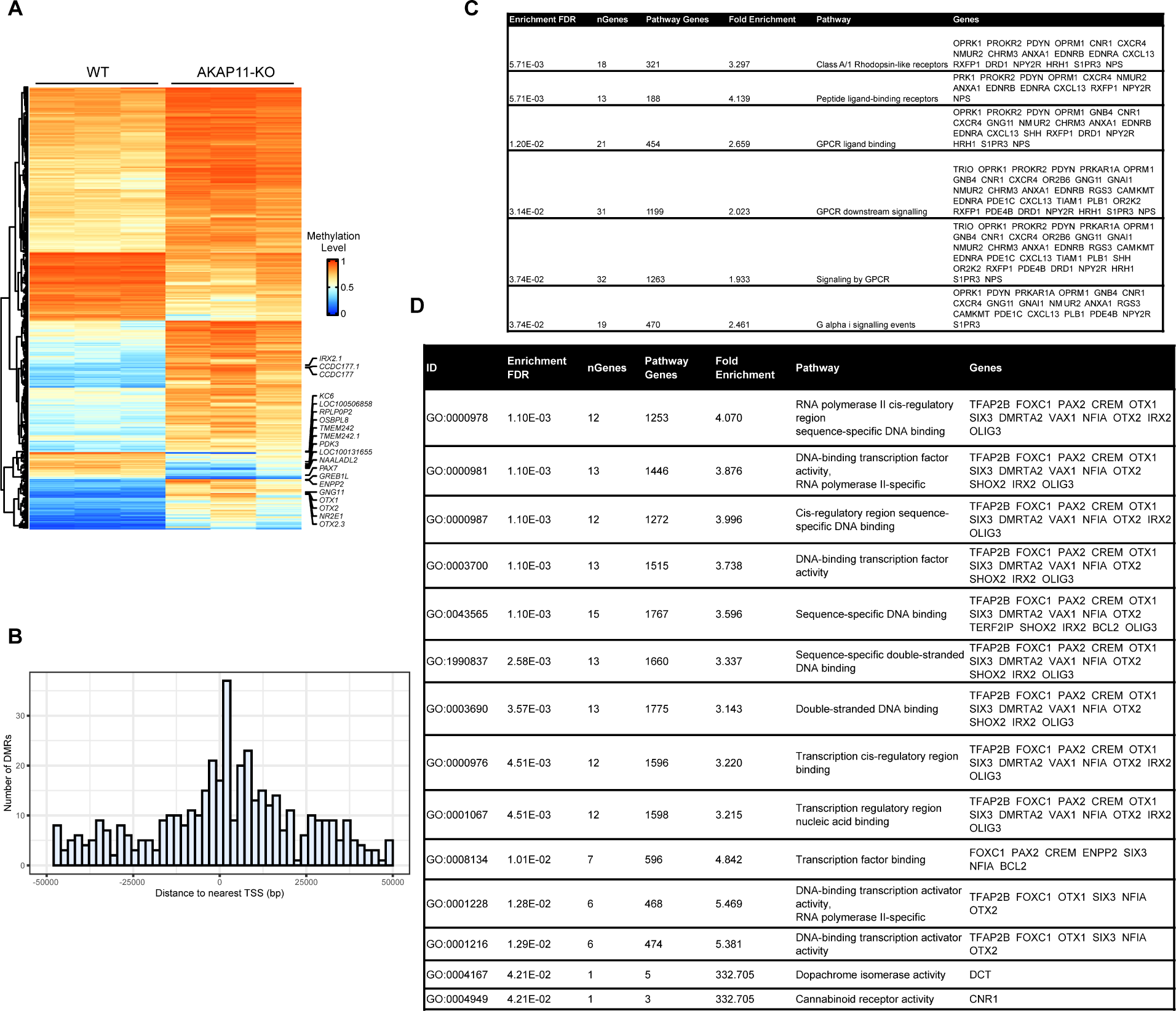
Heterozygous *AKAP11*-KO vs. WT DMR Characterization. **A)** Heatmap of patterns of methylation constructed using unsupervised hierarchical clustering of all DMRs identified (rows); columns represent the 3 WT replications and the 3 heterozygous *AKAP11*-KO clones. Orange and blue colors show the highest and lowest methylation levels, respectively. **B)** Histogram showing the locations of heterozygous *AKAP11*-KO vs. WT DMRs with respect to the closest TSS. TSS: transcription start site. **C)** Table of ORA of DMR-associated genes from heterozygous *AKAP11*-KO vs. WT comparison, using Curated.Reactome database in ShinyGO 0.77; pFDR < 0.05. The background genes used here were the same background gene list (23,590 genes that passed a low filter) from the RNA-Seq data. **D)** Table of top 20 GO molecular function terms (full list). ORA of genome-wide DMRs (regardless of their direction) that are associated with DEGs (regardless of their direction; at cutoff P-adj < 0.05 & |log2FC| > 0.25), regardless of their direction of change, using GO molecular function databases. Generated using ShinyGO 0.77; FDR < 0.05; top 10 pathways with redundancy removed. The background genes used here were the same background gene list (23,590 genes) obtained from our RNA-seq analysis.

**Supplementary Figure 4.**
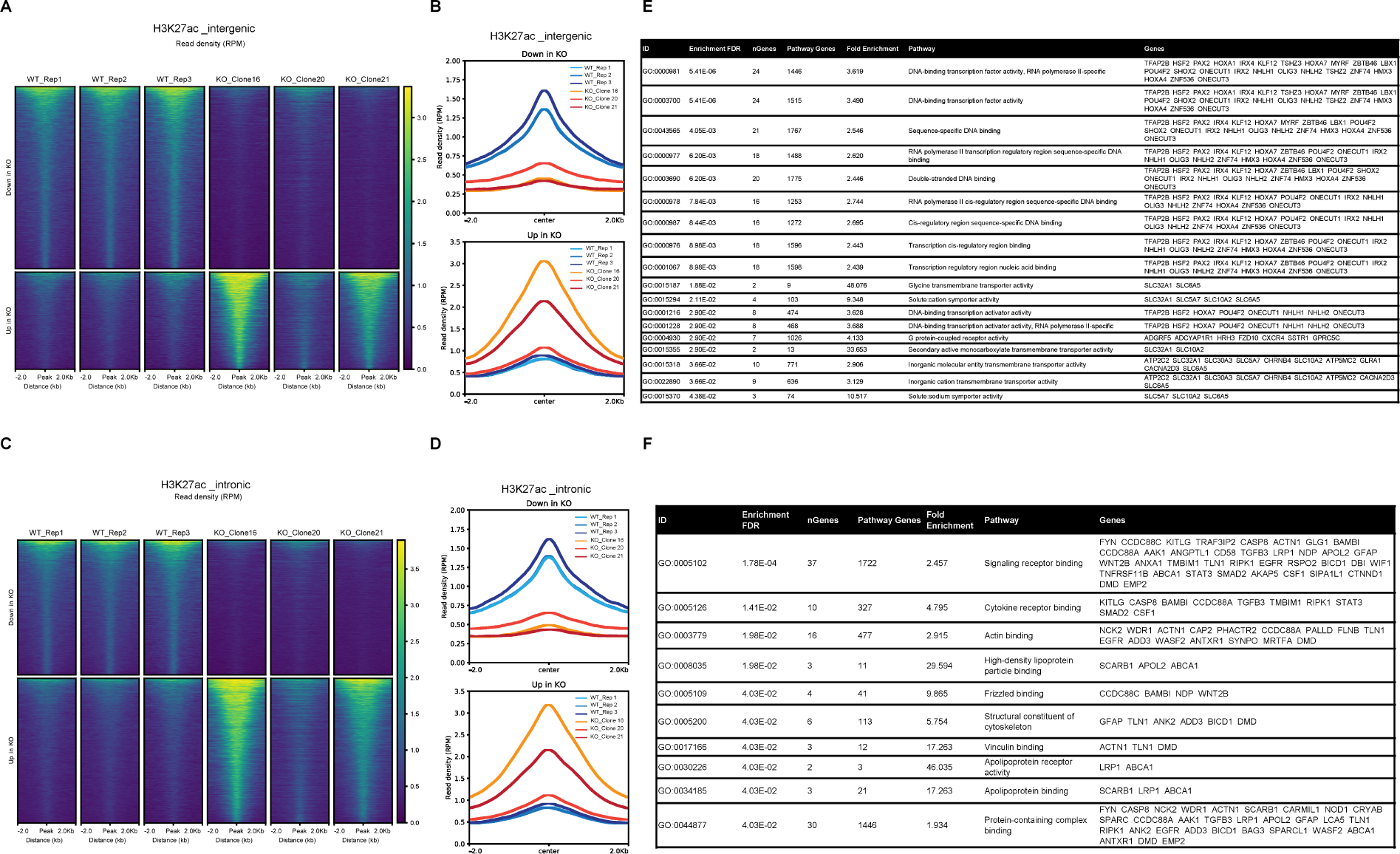
Intergenic and intronic differential H3K27ac peaks profiling and GO pathway analysis (GO-molecular function). **A)** Heatmap and **B)** average read density (RPM) profile aggregate plots of H3K27ac signals centered at differential intergenic peaks (center of enhancer ±2 kb flanking region), for heterozygous *AKAP11*-KO clones and WT replicates individually. **C)** Heatmap and **D)** average read density (RPM) profile plots of H3K27ac signals centered at differential intronic peaks (center of enhancer ±2 kb flanking region), for heterozygous *AKAP11*-KO clones and WT replicates individually. **E)** Results table of ORA of reduced intergenic H3K27ac peak intensities associated with decreased gene expression of target genes in heterozygous *AKAP11*-KO vs. WT (intergenic H3K27ac down, expression down: <u>group A</u>), using GO-Molecular Function database. Generated using ShinyGO 0.77; FDR < 0.09; top 20 pathways demonstrated. The background genes used here were the same background gene list (23,590 genes) obtained from our RNA-seq analysis. **F)** GO-Molecular Function ORA of increased intronic H3K27ac peak intensities associated with increased gene expression of target genes in heterozygous *AKAP11*-KO vs. WT (intronic H3K27ac up, expression up <u>group B</u>). Generated using ShinyGO 0.77; FDR < 0.05; top 20 pathways demonstrated. The background genes used here were the same background gene list (23,590 genes) obtained from our RNA-seq analysis.

### Supplementary Tables

**Supplementary Table 1**. Table of DEG results from RNA-seq data.

**Supplementary Table 2**. GSEA GO (molecular function) results from heterozygous *AKAP11*-KO vs. WT comparison. Padj<0.05.

**Supplementary Table 3. Table of** DMRs detected from WGBS data. DMRs were called using the function callDMR at default parameters with a minimum length of 50 base pairs and 3 CpG sites. Annotation determined using Homer (hg38).

**Supplementary Table 4.** DMR and DEG (p-adj < 0.05 and |log2FC| > 0.25) correlation plot gene list for heterozygous *AKAP11*-KO vs. WT.

**Supplementary Table 5.** Table of all genes from the correlation plots of differential H3K27ac peaks (p < 0.05 & |Log2FC| > 1) and differential gene expression (adj-p < 0.05 & |Log2FC| > 0.25) in intergenic as well as intronic regions.

**Supplementary Table 6.** Homer known motif enrichment results. Group A: Total Target Sequences = 188, Total Background Sequences = 49061. Group B: Total Target Sequences = 527, Total Background Sequences = 49150. p=1e-5.

**Supplementary Table 7**. Table of Differential H3K27ac (adj-P < 0.05 & |log2FC| > 1) and DMR Correlation in intergenic and intronic regions.

## ACKNOWLEDGMENTS

The work in G.R.’s lab is supported by the Canadian Institutes of Health Research and ERA PerMed. B.C.’s lab is supported by ERAPerMed PLOT-BD and a grant from the Fondation Bettencourt Schueller. We also acknowledge the financial support from the Brain & Behavior Research Foundation Young Investigator award to A.K. N.F. is supported by the studentship award Fonds de Recherche du Québec – Santé (FRQS). C.L. is supported by the CIHR Banting Fellowship. Computational analysis, data pre-processing, and adapted bioinformatics pipelines for analyses were performed by the Canadian Centre for Computational Genomics (C3G)-Montréal Node (Alain Pacis) using infrastructure provided by Compute Canada and Calcul Quebec. WGBS and ChIP-seq library preparation and sequencing were performed at McGill Genome Center platform. M.A. kindly donated the reprogrammed iPSCs to us.

## AUTHOR CONTRIBUTIONS

N.F. and C.L. conceived the study. N.F. designed, defined the scope, and carried out the laboratory experiments. A.K. and Y.L. gave guidance on the iPSC, NPC, and neuronal culture maintenance. N.F. designed and C3G (A.P.) carried out the computational analysis of the data with some guidance from N.F. and A.K. B.C. funded the next-generation sequencings and library preparations and G.R. lab funded the RNA-seq work and both provided guidance. M.A., P.D., G.R., B.C., C.L., and A.K. gave valuable scientific advice during the course of this study. N.F. wrote the manuscript.

## RESOURCE AVAILABILITY

### LEAD CONTACT

Additional information and requests for resources and reagents should be directed to and will be fulfilled by the Lead Contacts Drs. Rouleau, Khayachi, and Chaumette.

### MATERIALS AVAILABILITY

This study did not generate new unique reagents.

