## Supplementary Table 6 for "Transcriptomic and epigenomic consequences of heterozygous loss of function mutations in *AKAP11*, the first large-effect shared risk gene for bipolar disorder and schizophrenia"

### Homer Known Motif Enrichment Results (motif_INTERGENIC_H3K27ac-down_Expression-down)

[Homer *de novo* Motif Results](#)
[Gene Ontology Enrichment Results](#)
[Known Motif Enrichment Results (txt file)](#)
Total Target Sequences = 188, Total Background Sequences = 49061

| Rank | Motif | Name | P-value | log P-pvalue | q-value (Benjamini) | # Target Sequences with Motif | % of Targets Sequences with Motif | # Background Sequences with Motif | % of Background Sequences with Motif | Motif File | SVG |
|---|---|---|---|---|---|---|---|---|---|---|---|
| 1 | 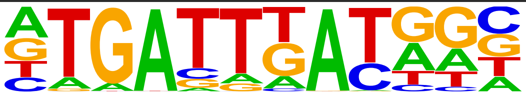 | PBX2(Homeobox)/K562-PBX2-ChIP-Seq(Encode)/Homer | 1e-3 | -8.211e+00 | 0.1195 | 29.0 | 15.43% | 3777.7 | 7.70% | motif file (matrix) | svg |
| 2 | 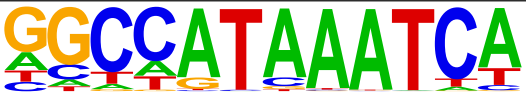 | Hoxc9(Homeobox)/Ainv15-Hoxc9-ChIP-Seq(GSE21812)/Homer | 1e-2 | -6.181e+00 | 0.4549 | 20.0 | 10.64% | 2556.4 | 5.21% | motif file (matrix) | svg |
| 3 | 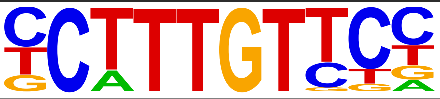 | Sox4(HMG)/proB-Sox4-ChIP-Seq(GSE50066)/Homer | 1e-2 | -5.569e+00 | 0.5596 | 25.0 | 13.30% | 3675.3 | 7.49% | motif file (matrix) | svg |
| 4 | 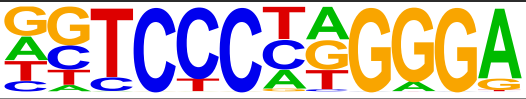 | EBF(EBF)/proBcell-EBF-ChIP-Seq(GSE21978)/Homer | 1e-2 | -5.127e+00 | 0.6526 | 10.0 | 5.32% | 1009.9 | 2.06% | motif file (matrix) | svg |
| 5 | 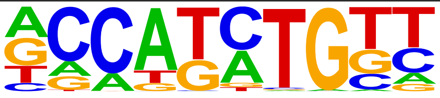 | Olig2(bHLH)/Neuron-Olig2-ChIP-Seq(GSE30882)/Homer | 1e-2 | -5.095e+00 | 0.6526 | 50.0 | 26.60% | 9277.3 | 18.91% | motif file (matrix) | svg |
| 6 | 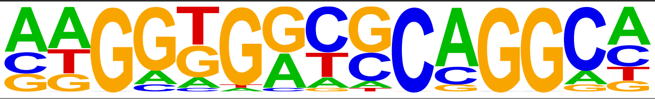 | ZNF165(Zf)/WHIM12-ZNF165-ChIP-Seq(GSE65937)/Homer | 1e-2 | -5.015e+00 | 0.6526 | 8.0 | 4.26% | 713.3 | 1.45% | motif file (matrix) | svg |
| 7 | 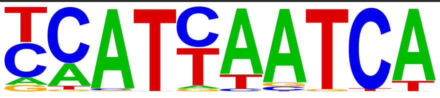 | Pdx1(Homeobox)/Islet-Pdx1-ChIP-Seq(SRA008281)/Homer | 1e-2 | -4.754e+00 | 0.6526 | 29.0 | 15.43% | 4764.5 | 9.71% | motif file (matrix) | svg |

### Homer Known Motif Enrichment Results (motif_INTRONIC_H3K27ac-up_Expression-up)

Homer *de novo* Motif Results
Gene Ontology Enrichment Results
Known Motif Enrichment Results (txt file)
Total Target Sequences = 527, Total Background Sequences = 49150

| Rank | Motif | Name | P-value | log P-pvalue | q-value (Benjamini) | # Target Sequences with Motif | % of Targets Sequences with Motif | # Background Sequences with Motif | % of Background Sequences with Motif | Motif File | SVG |
|---|---|---|---|---|---|---|---|---|---|---|---|
| 1 | 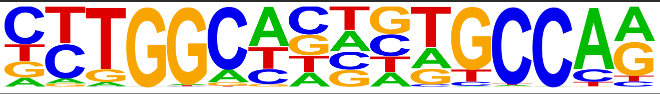 | NF1(CTF)/LNCAP-NF1-ChIP-Seq(Unpublished)/Homer | 1e-5 | -1.188e+01 | 0.0030 | 40.0 | 7.59% | 1735.0 | 3.53% | motif file (matrix) | svg |
| 2 | 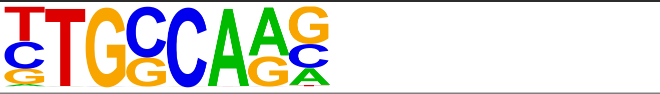 | NF1-halfsite(CTF)/LNCaP-NF1-ChIP-Seq(Unpublished)/Homer | 1e-4 | -9.617e+00 | 0.0147 | 124.0 | 23.53% | 8328.6 | 16.93% | motif file (matrix) | svg |
| 3 | 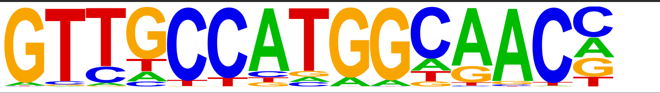 | Rfx2(HTH)/LoVo-RFX2-ChIP-Seq(GSE49402)/Homer | 1e-3 | -8.886e+00 | 0.0203 | 10.0 | 1.90% | 215.3 | 0.44% | motif file (matrix) | svg |
| 4 | 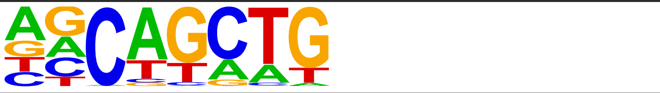 | SCL(bHLH)/HPC7-Scl-ChIP-Seq(GSE13511)/Homer | 1e-3 | -8.881e+00 | 0.0203 | 268.0 | 50.85% | 21095.1 | 42.88% | motif file (matrix) | svg |
| 5 | 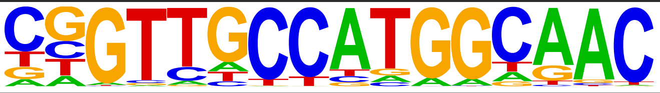 | RFX(HTH)/K562-RFX3-ChIP-Seq(SRA012198)/Homer | 1e-3 | -8.694e+00 | 0.0203 | 9.0 | 1.71% | 179.5 | 0.36% | motif file (matrix) | svg |
| 6 | 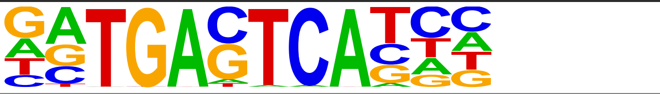 | Jun-AP1(bZIP)/K562-cJun-ChIP-Seq(GSE31477)/Homer | 1e-3 | -8.636e+00 | 0.0203 | 21.0 | 3.98% | 792.2 | 1.61% | motif file (matrix) | svg |
| 7 | 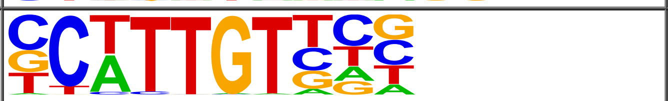 | Sox10(HMG)/SciaticNerve-Sox3-ChIP-Seq(GSE35132)/Homer | 1e-3 | -8.392e+00 | 0.0203 | 113.0 | 21.44% | 7663.3 | 15.58% | motif file (matrix) | svg |
| 8 | 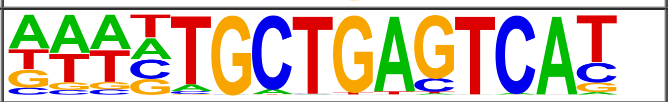 | Bach1(bZIP)/K562-Bach1-ChIP-Seq(GSE31477)/Homer | 1e-3 | -8.147e+00 | 0.0203 | 9.0 | 1.71% | 193.9 | 0.39% | motif file (matrix) | svg |
| 9 | 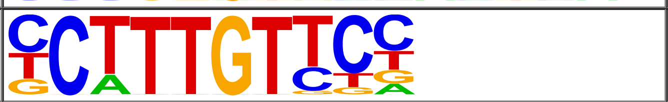 | Sox4(HMG)/proB-Sox4-ChIP-Seq(GSE50066)/Homer | 1e-3 | -8.105e+00 | 0.0203 | 66.0 | 12.52% | 3979.0 | 8.09% | motif file (matrix) | svg |
| 10 | 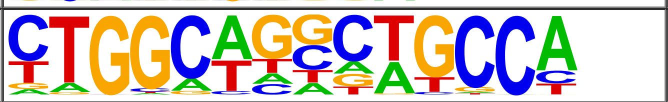 | Tlx?(NR)/NPC-H3K4me1-ChIP-Seq(GSE16256)/Homer | 1e-3 | -7.986e+00 | 0.0203 | 36.0 | 6.83% | 1805.8 | 3.67% | motif file (matrix) | svg |
| 11 | 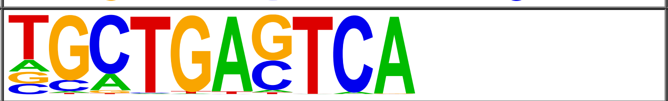 | Bach2(bZIP)/OCILy7-Bach2-ChIP-Seq(GSE44420)/Homer | 1e-3 | -7.745e+00 | 0.0203 | 18.0 | 3.42% | 670.5 | 1.36% | motif file (matrix) | svg |
| 12 | 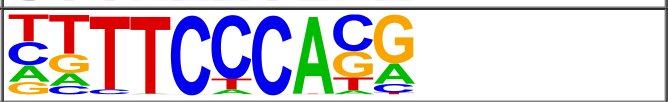 | Rbpj1(?)/Panc1-Rbpj1-ChIP-Seq(GSE47459)/Homer | 1e-3 | -7.412e+00 | 0.0222 | 102.0 | 19.35% | 6958.6 | 14.15% | motif file (matrix) | svg |
| 13 | 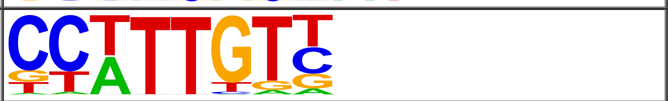 | Sox3(HMG)/NPC-Sox3-ChIP-Seq(GSE33059)/Homer | 1e-3 | -7.365e+00 | 0.0222 | 121.0 | 22.96% | 8542.3 | 17.37% | motif file (matrix) | svg |
| 14 | 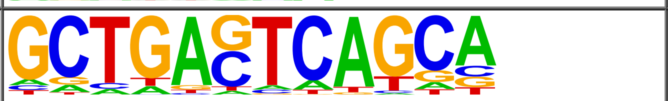 | MafK(bZIP)/C2C12-MafK-ChIP-Seq(GSE36030)/Homer | 1e-3 | -7.194e+00 | 0.0236 | 21.0 | 3.98% | 886.3 | 1.80% | motif file (matrix) | svg |
| 15 | 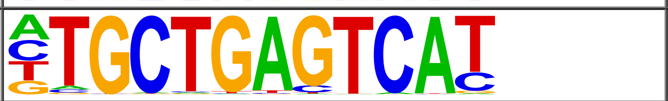 | Nrf2(bZIP)/Lymphoblast-Nrf2-ChIP-Seq(GSE37589)/Homer | 1e-3 | -6.937e+00 | 0.0285 | 8.0 | 1.52% | 184.9 | 0.38% | motif file (matrix) | svg |
| 16 | 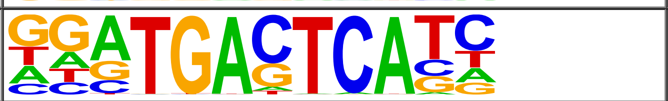 | Fra2(bZIP)/Striatum-Fra2-ChIP-Seq(GSE43429)/Homer | 1e-2 | -6.902e+00 | 0.0285 | 33.0 | 6.26% | 1708.0 | 3.47% | motif file (matrix) | svg |
| 17 | 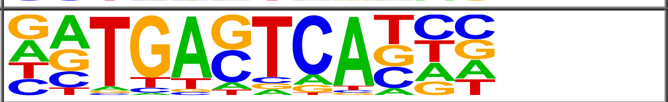 | Fosl2(bZIP)/3T3L1-Fosl2-ChIP-Seq(GSE56872)/Homer | 1e-2 | -6.660e+00 | 0.0332 | 24.0 | 4.55% | 1120.3 | 2.28% | motif file (matrix) | svg |
| 18 | 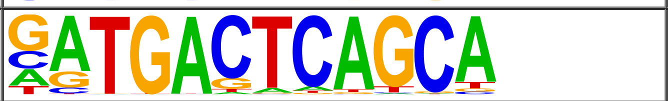 | NF-E2(bZIP)/K562-NFE2-ChIP-Seq(GSE31477)/Homer | 1e-2 | -6.637e+00 | 0.0332 | 8.0 | 1.52% | 193.4 | 0.39% | motif file (matrix) | svg |
| 19 | 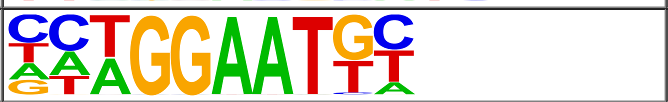 | TEAD(TEA)/Fibroblast-PU.1-ChIP-Seq(Unpublished)/Homer | 1e-2 | -6.580e+00 | 0.0332 | 56.0 | 10.63% | 3445.7 | 7.00% | motif file (matrix) | svg |
| 20 | 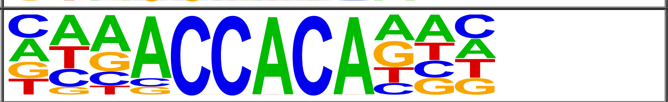 | RUNX2(Runt)/PCa-RUNX2-ChIP-Seq(GSE33889)/Homer | 1e-2 | -6.378e+00 | 0.0374 | 60.0 | 11.39% | 3787.0 | 7.70% | motif file (matrix) | svg |
| 21 | 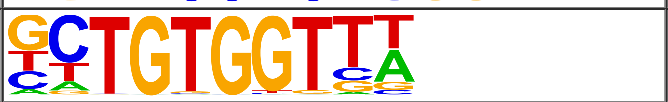 | RUNX-AML(Runt)/CD4+-PolII-ChIP-Seq(Barski_et_al.)/Homer | 1e-2 | -6.206e+00 | 0.0423 | 53.0 | 10.06% | 3273.2 | 6.65% | motif file (matrix) | svg |
| 22 | 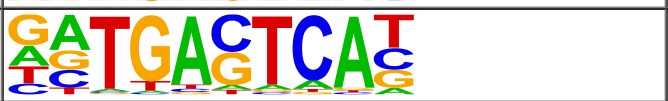 | JunB(bZIP)/DendriticCells-Junb-ChIP-Seq(GSE36099)/Homer | 1e-2 | -5.959e+00 | 0.0517 | 36.0 | 6.83% | 2031.3 | 4.13% | motif file (matrix) | svg |
| 23 | 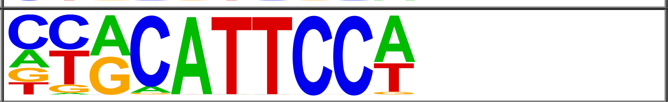 | TEAD1(TEAD)/HepG2-TEAD1-ChIP-Seq(Encode)/Homer | 1e-2 | -5.890e+00 | 0.0529 | 70.0 | 13.28% | 4666.7 | 9.49% | motif file (matrix) | svg |
| 24 | 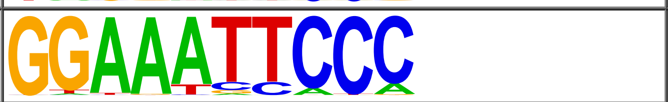 | NFkB-p65-Rel(RHD)/ThioMac-LPS-Expression(GSE23622)/Homer | 1e-2 | -5.837e+00 | 0.0535 | 8.0 | 1.52% | 220.2 | 0.45% | motif file (matrix) | svg |
| 25 | 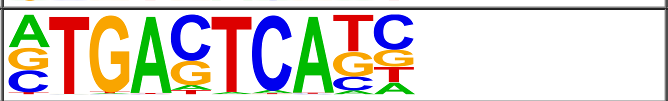 | AP-1(bZIP)/ThioMac-PU.1-ChIP-Seq(GSE21512)/Homer | 1e-2 | -5.725e+00 | 0.0574 | 47.0 | 8.92% | 2888.5 | 5.87% | motif file (matrix) | svg |
| 26 | 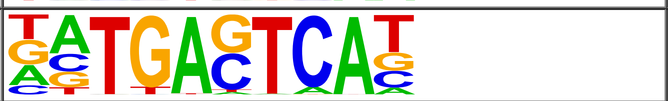 | BATF(bZIP)/Th17-BATF-ChIP-Seq(GSE39756)/Homer | 1e-2 | -5.719e+00 | 0.0574 | 42.0 | 7.97% | 2509.4 | 5.10% | motif file (matrix) | svg |
| 27 | 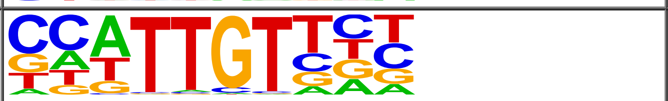 | Sox6(HMG)/Myotubes-Sox6-ChIP-Seq(GSE32627)/Homer | 1e-2 | -5.718e+00 | 0.0574 | 108.0 | 20.49% | 7839.8 | 15.94% | motif file (matrix) | svg |
| 28 | 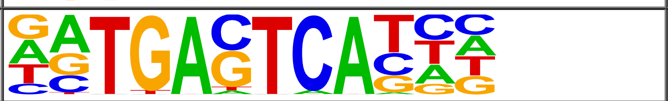 | Atf3(bZIP)/GBM-ATF3-ChIP-Seq(GSE33912)/Homer | 1e-2 | -5.691e+00 | 0.0574 | 42.0 | 7.97% | 2513.6 | 5.11% | motif file (matrix) | svg |

| | | | | | | | | | | | |
|---|---|---|---|---|---|---|---|---|---|---|---|
| 29 | 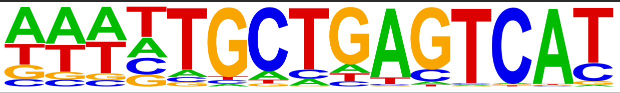 | NFE2L2(bZIP)/HepG2-NFE2L2-ChIP-Seq(Encode)/Homer | 1e-2 | -5.552e+00 | 0.0588 | 7.0 | 1.33% | 182.6 | 0.37% | motif file (matrix) | svg |
| 30 | 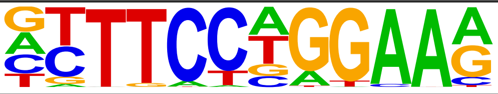 | STAT4(Stat)/CD4-Stat4-ChIP-Seq(GSE22104)/Homer | 1e-2 | -5.206e+00 | 0.0805 | 67.0 | 12.71% | 4561.1 | 9.27% | motif file (matrix) | svg |
| 31 | 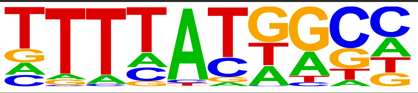 | Hoxa11(Homeobox)/ChickenMSG-Hoxa11.Flag-ChIP-Seq(GSE86088)/Homer | 1e-2 | -5.141e+00 | 0.0830 | 166.0 | 31.50% | 13028.7 | 26.49% | motif file (matrix) | svg |
| 32 | 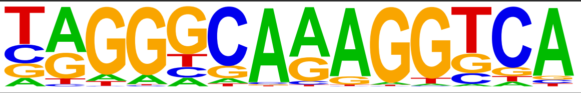 | RXR(NR),DR1/3T3L1-RXR-ChIP-Seq(GSE13511)/Homer | 1e-2 | -5.088e+00 | 0.0848 | 72.0 | 13.66% | 4993.8 | 10.15% | motif file (matrix) | svg |
| 33 | 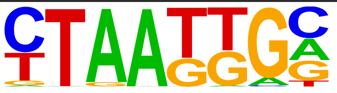 | Isl1(Homeobox)/Neuron-Isl1-ChIP-Seq(GSE31456)/Homer | 1e-2 | -5.036e+00 | 0.0867 | 130.0 | 24.67% | 9909.0 | 20.14% | motif file (matrix) | svg |
| 34 | 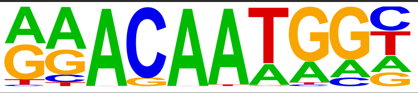 | Sox15(HMG)/CPA-Sox15-ChIP-Seq(GSE62909)/Homer | 1e-2 | -4.992e+00 | 0.0879 | 79.0 | 14.99% | 5591.7 | 11.37% | motif file (matrix) | svg |
| 35 | 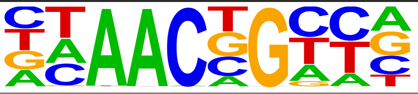 | BMYB(HTH)/Hela-BMYB-ChIP-Seq(GSE27030)/Homer | 1e-2 | -4.988e+00 | 0.0879 | 92.0 | 17.46% | 6676.1 | 13.57% | motif file (matrix) | svg |
| 36 | 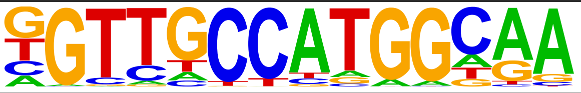 | Rfx1(HTH)/NPC-H3K4me1-ChIP-Seq(GSE16256)/Homer | 1e-2 | -4.928e+00 | 0.0885 | 15.0 | 2.85% | 676.9 | 1.38% | motif file (matrix) | svg |
| 37 | 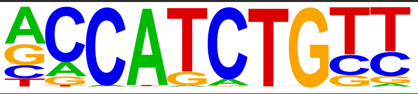 | NeuroG2(bHLH)/Fibroblast-NeuroG2-ChIP-Seq(GSE75910)/Homer | 1e-2 | -4.886e+00 | 0.0898 | 97.0 | 18.41% | 7124.1 | 14.48% | motif file (matrix) | svg |
| 38 | 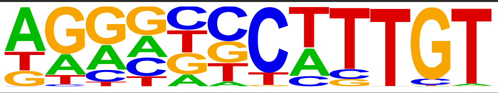 | Sox9(HMG)/Limb-SOX9-ChIP-Seq(GSE73225)/Homer | 1e-2 | -4.721e+00 | 0.1032 | 55.0 | 10.44% | 3692.1 | 7.51% | motif file (matrix) | svg |
| 39 | 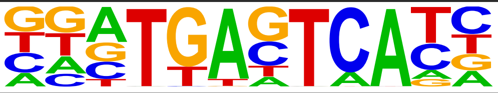 | Fos(bZIP)/TSC-Fos-ChIP-Seq(GSE110950)/Homer | 1e-2 | -4.719e+00 | 0.1032 | 36.0 | 6.83% | 2203.8 | 4.48% | motif file (matrix) | svg |
| 40 | 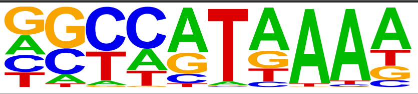 | Hoxd11(Homeobox)/ChickenMSG-Hoxd11.Flag-ChIP-Seq(GSE86088)/Homer | 1e-2 | -4.687e+00 | 0.1032 | 165.0 | 31.31% | 13092.6 | 26.62% | motif file (matrix) | svg |
| 41 | 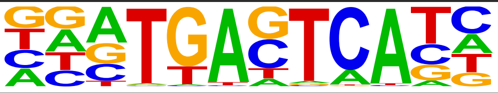 | Fra1(bZIP)/BT549-Fra1-ChIP-Seq(GSE46166)/Homer | 1e-2 | -4.672e+00 | 0.1032 | 34.0 | 6.45% | 2058.1 | 4.18% | motif file (matrix) | svg |
| 42 | 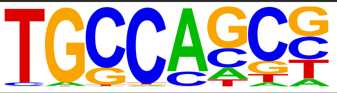 | HIC1(Zf)/Treg-ZBTB29-ChIP-Seq(GSE99889)/Homer | 1e-2 | -4.668e+00 | 0.1032 | 133.0 | 25.24% | 10282.7 | 20.90% | motif file (matrix) | svg |
| 43 | 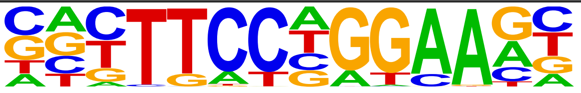 | Stat3+il21(Stat)/CD4-Stat3-ChIP-Seq(GSE19198)/Homer | 1e-2 | -4.643e+00 | 0.1032 | 47.0 | 8.92% | 3070.1 | 6.24% | motif file (matrix) | svg |
| 44 | 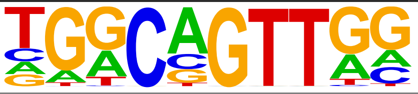 | AMYB(HTH)/Testes-AMYB-ChIP-Seq(GSE44588)/Homer | 1e-2 | -4.617e+00 | 0.1032 | 91.0 | 17.27% | 6688.7 | 13.60% | motif file (matrix) | svg |
| 45 | 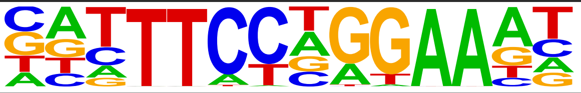 | STAT1(Stat)/HelaS3-STAT1-ChIP-Seq(GSE12782)/Homer | 1e-2 | -4.608e+00 | 0.1032 | 24.0 | 4.55% | 1326.2 | 2.70% | motif file (matrix) | svg |
